# Model substrate particles uncover dynamics of microbial communities associated with particulate organic matter decomposition in soil

**DOI:** 10.1101/2025.06.27.661937

**Authors:** Eva Simon, Lauren V. Alteio, Alexander König, Bruna Imai, Julia Horak, Julia Wiesenbauer, Joana Séneca, Bela Hausmann, Marc Mussmann, Barbara Kitzler, Christina Kaiser

## Abstract

Soil organic matter is the largest terrestrial reservoir of organic carbon. Its particulate fraction, particulate organic matter (POM), serves as a resource and surface for microbial colonization. Degradation of complex biopolymers like cellulose and chitin requires extracellular enzymes produced by phylogenetically diverse microbes. Despite their importance for carbon cycling, the structure and spatio-temporal dynamics of POM-associated microbial communities in soil and how specific substrates influence them remain poorly understood. This study investigated whether microbial communities associated with POM change in composition and richness over time and whether chitin and cellulose select for distinct fungal and bacterial taxa. We incubated self-manufactured, millimetre-sized model substrate particles containing chitin or cellulose in soil under laboratory and field conditions. We assessed particle-associated communities at multiple time points over a 50-day-long incubation in the lab and after 47 days in the field. Our results show that community structure and temporal dynamics of particle-associated microbial communities were substrate-specific. While microbial biomass increased on both particle types, chitin-associated microbial communities exhibited stronger temporal changes. Communities on chitin and cellulose particles were enriched in specific bacterial and fungal genera compared to communities in the surrounding soil. We demonstrate that microbial communities associated with model chitin particles underwent notable temporal changes, including decreased microbial richness and shifts in community composition over the incubation period. This study shows the potential of model particles to advance our understanding of particle- and substrate-associated microbial communities in soil.

## Introduction

Soil organic matter is the largest terrestrial carbon reservoir [1, 2]. Conceptually, soil organic matter can be divided into mineral-associated organic matter and particulate organic matter (POM) [3]. POM serves as a surface for colonization [4] and particles are considered hotspots for microbial activity [5] as they represent spatially isolated, nutrient-rich microhabitats in the otherwise nutrient-poor soil environment.

Microbes decompose POM, as they can only take up small molecules (typically less than 600 Da) [1, 6]. To acquire resources, they produce and excrete extracellular enzymes [7, 8], which break down large molecules into subunits and ultimately monomers that can be taken up [9, 10]. By decomposing organic matter via extracellular enzymes, heterotrophic microorganisms mediate the recycling of carbon and nutrients [9, 11–13]. The complete breakdown of biopolymers requires multiple enzymes that act in synergy [14–16]. Extracellular enzymes are substrate-specific [7] and metabolically costly to produce [8, 16]. Genetic potential to produce extracellular enzymes differs between microbial taxa [6, 17–19]. Consequently, phylogenetically diverse microbial communities are necessary to complete biopolymer degradation.

Cellulose and chitin are the two most abundant polymers in nature [18, 20, 21]. Both must be enzymatically degraded prior to microbial uptake. While the potential to degrade these polysaccharides is widespread, the structure of degrader communities is driven by environmental conditions and substrate availability [18, 22]. Amongst fungi, the ability to degrade cellulose is more widespread than the ability to degrade chitin [12].

Spatial proximity of microbial cells to enzymatically degraded substrates is advantageous to enhance the return on investment. Microbial attachment to POM and formation of spatially structured communities should minimize losses of enzymatic breakdown products to the surrounding environment through diffusion. The presence of both bacteria and fungi has been detected on POM in soil. More specifically, it has been shown that bacteria occur in biofilms on the surface of POM and fungal hyphae pervade bacterial biofilms [5]. However, little is known about microbial communities associated with POM, despite the relevance of POM-degrading communities for carbon and nutrient cycling. We lack knowledge about the assembly and temporal dynamics of microbial communities on POM, and whether specific substrates select distinct microbial communities and particular microbial taxa in soil.

This knowledge gap is partly due to the complex nature of soil, making it challenging to study microbial communities directly associated with POM in soil. Soils are opaque and characterized by high substrate heterogeneity [4]. Organic matter exhibits diverse chemical composition and is spatially heterogeneously distributed throughout the soil matrix [3, 23–26]. Investigation of microbial communities associated with natural POM requires extraction with chemical agents, which may alter associated microbial communities [27, 28], possibly resulting in erroneous findings. Other studies investigated substrate-associated communities using isotopic labelling or metagenomics [29, 30], however, community dynamics at the millimetre-spatial scale remain poorly understood.

In this study, we assess how the complex substrates, chitin and cellulose, drive successional changes in microbial community assembly. Specifically, we address microbial colonization and assembly at the particle interface in the heterogeneous soil environment using substrate-containing model particles. We hypothesized that (i) microbial colonization of particles is highly substrate-specific due to the degree of substrate complexity. Additionally, (ii) changes in microbial communities are more pronounced at the particle-scale than at the soil-scale due to the relevance of the micro-scale for microbial activity and interactions. Finally, (iii) substrate-driven changes in community composition observed under controlled laboratory conditions are reproducible in the field due to the fundamental influence of substrate in driving community assembly. We applied self-manufactured model particles containing either chitin or cellulose to study microbial communities at the particle interface in soil. While model particles differ from natural POM in their physical and chemical properties, they provide a simple and tractable experimental system which facilitates the study of microbial communities associated with complex substrates in particulate form. The promising applications of model substrate particles in microbial ecology were recently highlighted [31]. Model particles have been used to study microbial community assembly and dynamics on POM surfaces in aquatic systems [32–35], but, to our knowledge, not in soils. We incubated model substrate particles in forest soil and examined associated microbial communities associated with particles and surrounding soil through a series of sampling time points over

50 days in the lab. In parallel, we incubated particles in soil in the field for 47 days to investigate microbial community assembly under more natural conditions. At harvest time points, we performed amplicon sequencing of model particle fractions and the surrounding soil and assessed taxa richness and community composition. In addition, we determined microbial biomass on particles and in soil via droplet digital PCR.

## Materials and Methods

### Model particle production

We manufactured three types of model particles: (i) particles containing chitin (“model chitin particles”), (ii) cellulose (“model cellulose particles”), and (iii) particles containing neither chitin nor cellulose (“no substrate”). We used Gelrite**®** gellan gum (Carl Roth, Karlsruhe, DE) to produce model particles because of its physical properties (e.g., rapid and stable gelling with ionic cross-linkers, high transparency) and its resistance to enzymatic degradation, making it a relatively biologically inert scaffold for embedding substrates compared to other frequently used gelling agents, like alginate and agarose [36, 37]. Following a protocol by Mussmann [38], we dissolved 1.5 g of Gelrite® gellan gum in 100 mL of MilliQ water while stirring. To facilitate polymer cross-linking, CaCl_2_ (Sigma-Aldrich, Vienna, AT) was added at a final concentration of 8 mM. Additionally, we added 5 mL of 10% w/v iron (II, III) oxide (Sigma-Aldrich, Vienna, AT) to facilitate magnetic separation in downstream washing steps. As the only modification between particle types, we suspended 1.4 g of chitin from shrimp shells (Sigma-Aldrich, Vienna, AT) or cellulose powder (Sigma-Aldrich, Vienna, AT) in the gellan gum solution and applied an additional 0.5 g of respective substrate to the bottom and top of the poured medium in a petri plate prior to solidification. No substrate was supplemented to the “no substrate” control particles. Model particles were cylindrical and approximately 3 x 2 mm. Particles were UV-sterilized and stored in 20% ethanol at 4°C until use.

### Soil collection and processing

Soil was collected in a beech-dominated temperate forest at the long-term ecological research site (LTER) in Klausen-Leopoldsdorf, Austria (48.11925 N, 16.04343 E, 580 m altitude) on 26 April 2021.This soil is classified as a dystric cambisol over sandstone with a loam-loamy clay texture [39–41]. The litter layer was removed prior to sampling, where soil cores were taken from the upper 5 cm of mineral soil at five established 5 x 5 m field plots [42] using a 5 x 8 cm metal corer. For each plot, two soil cores were pooled and homogenized into a single representative sample using a 2 mm mesh sieve, producing five composite soil samples. We recorded a gravimetric soil water content of 0.52 ± 0.13 g water g^-1^ dry soil (median ± standard deviation) and a pH of 4.85 ± 0.18 for sieved soil samples used for lab and field incubations.

### Set-up of laboratory and field incubation

We incubated model particles in sieved forest soil in the laboratory and the field. For the lab incubation, we prepared mesocosms using 50 mL plastic vials (Greiner, Kremsmuenster, AT) by cutting them at 5 cm height with their cap serving as the bottom of the mesocosm. For the field incubation, we sewed mesh bags from polyester screen-printing fabric with a mesh width of 40 µm (Siebdruckversand, Magdeburg, DE) to allow fungal hyphal ingrowth but exclude roots, mimicking lab conditions [43, 44].

We set up four incubation treatments: i) 2-mm-sieved soil plus model chitin particles, ii) soil plus model cellulose particles, iii) soil plus “no substrate” model particles containing neither chitin nor cellulose, and iv) soil without model particles (Fig. 1). We rinsed model particles, which had been stored in 20% ethanol, three times in 15 mL pure MilliQ water to remove residual ethanol. We added 1 g of model particles to 15 g of sieved soil to individual plastic bags to homogenize and distribute model particles evenly. Additionally, 15 g sieved soil which was incubated without model particles was homogenized. Soil (and model particles) was filled in lab mesocosms and field mesh bags (28 April 2021). Mesocosms were compacted to field density (1.0 g cm^-3^). Mesocosms were covered with parafilm during incubation to prevent desiccation. Mesh bags were sealed with a heat-sealer (Mercier Corporation, Taiwan) and measured 7 x 10 cm in size. In total, we set up 80 mesocosms and 20 mesh bags (Fig. 1).

**Fig. 1.**
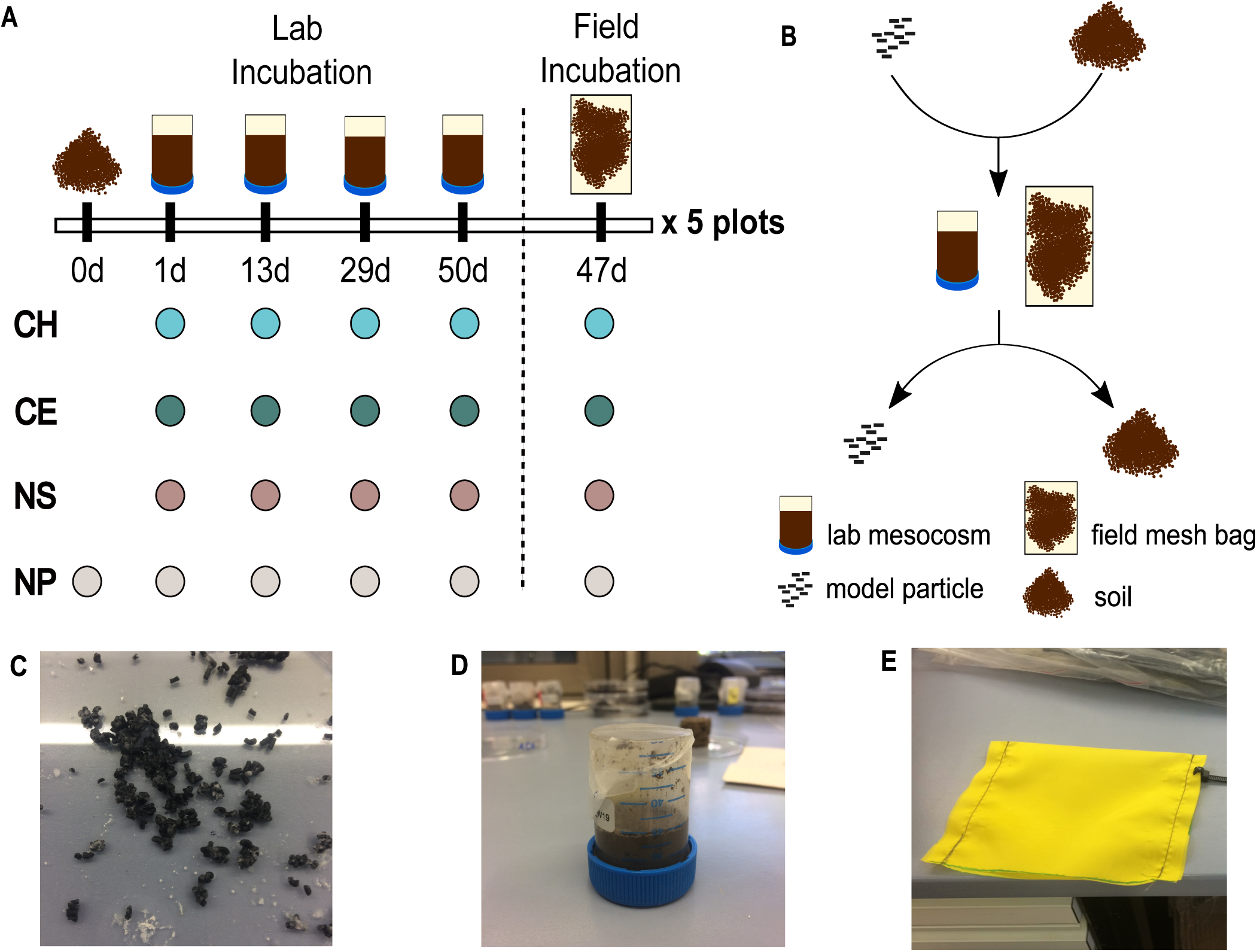
Schematic of the experimental set-up. (**A**) We set up four incubation treatments, incubating (i) self-manufactured model chitin (CH), (ii) cellulose (CE), (iii) “no substrate” (NS) particles and (iv) no model particles in 2-mm-sieved soil. We investigated five biological replicates by using soil sampled from five field plots. We set up four technical replicates per treatment and plot for the lab incubation, which we harvested at four time points, after one, 13, 29 or 50 days of incubation in the laboratory, and one technical replicate per treatment and plot for incubation in the field which we harvested after 47 days. (**B**) We mixed 1 g of model particles and 15 g 2-mm-sieved soil and filled it in lab mesocosms that were produced from 50 mL Greiner tubes or mesh bags for the field. Upon harvest of mesocosms and mesh bags, we manually separated model particles from the soil for subsequent analysis. (**C**) Example of self-manufactured model chitin substrate particles prior to incubation. (**D**) Laboratory mesocosm filled with soil and model particles. (**E**) Mesh bag used for the field incubation.

Mesocosms were incubated in an incubator at 11°C ± 2°C, matching the soil temperature at 5 cm soil depth on the day of soil sampling. Filled mesh bags were kept in closed plastic bags at 11°C in the lab incubator until they were buried in the field at a depth of 5 cm (30 April 2021). Potential moisture loss was assessed by weighing mesocosms weekly and adjusted by adding MilliQ water if weight deviated by ≥ 2%.

### Destructive harvest

Lab mesocosms were destructively harvested one, 13, 29, and 50 days after the start of the lab incubation (start on 28 April; harvest on 29 April, 11 May, 27 May, 17 June 2021). Mesh bags were destructively harvested after 47 days of incubation in the field (filled on 28 April; buried on 30 April; harvest on 14 June 2021). Upon destructive harvest, we extracted the mesocosms from the vials and separated model particles from soil using tweezers. Separated particles were rinsed three times with MilliQ water to remove lightly adhering soil and were (i) weighed for DNA extraction and (ii) fixed with ethanol for downstream microscopic analysis.

### DNA extraction and sequencing

DNA extraction, library preparation, and amplicon sequencing were conducted as in Simon et al (2024) [42]. In short, we extracted DNA from a minimum of 20 model particles (150 mg) and 250 mg soil with the Qiagen PowerSoil kit (Qiagen, Valencia, CA, USA), following the manufacturer’s protocol with exception of an added bead beating step (30 sec). For amplicon sequencing, we followed a two-step PCR barcoding approach [45] to amplify the V4 region of the 16S rRNA genes targeting bacteria and the fungal internal transcribed spacer region 2 (ITS2) of the nuclear rRNA operon. We used primers 515F [46] and 806R [47] for 16S rRNA genes. For the ITS2 region, a nested approach [48] was chosen, generating a PCR product using primers ITSOF-T [49, 50] and ITS4 [51], which was used as a template for a second PCR using primers gITS7 [52] and ITS4. Amplicons were sequenced on a MiSeq platform (Illumina, V3 chemistry, 600 cycles) at the Joint Microbiome Facility (JMF) of the Medical University of Vienna and the University of Vienna under project number JMF-2106-12. Raw data processing was performed as described previously [45].

### Droplet digital PCR (ddPCR)

We quantified the number of copies of the V4 hypervariable region of the 16S rRNA gene and of the fungal ITS region of model particles and in soils with droplet digital PCR (ddPCR). Gene copy numbers can be used as a proxy for microbial biomass [53]. Prior to performing ddPCR, we measured DNA concentration of all DNA extracts using PicoGreen (Quant-iT PicoGreen dsDNA Assay Kit; Thermo Fisher Scientific, Vienna, AT). Extracts were re-measured using a Qubit high-sensitivity dsDNA assay (Thermo Fisher Scientific, Vienna, AT) if the concentration was under the detection limit. For 16S ddPCR, DNA extracts were diluted to 0.05 ng µL^-1^ and for ITS ddPCR to 0.25 ng µL^-1^. 2 µL of diluted DNA template were used as input for each ddPCR assay. Gene copy numbers per µL DNA extract were assessed for a single replicate of each sample with the QX200 Droplet Digital PCR (Bio-Rad) and analysed with QuantaSoft software (Bio-Rad, Vienna, AT).

### Particle fixation and fluorescence microscopy

A fraction of incubated model particles (∼ 12 model particles) were fixed with 96% ethanol in 2 mL Eppendorf tubes (Sigma-Aldrich, Vienna, AT) and stored overnight at 4°C. Supernatant was removed using magnetic pulldown and model particles were resuspended in 1.5 mL of 1X PBS to wash a total of three times. Ethanol-fixed samples were suspended in 1.5 mL of cold 96% ethanol. Following fixation, samples were stored at −20°C. Selected chitin particles were stained with SYBR Green I (Invitrogen, Thermo Fisher Scientific, Vienna, AT) and analysed using a Leica confocal laser scanning microscope, coupled with a Thunder imaging system (Leica, Vienna, AT).

### Data analyses

Amplicon sequence variants (ASVs) were inferred using the DADA2 R package v1.20 (https://www.ncbi.nlm.nih.gov/pubmed/27214047) applying the recommended workflow (https://f1000research.com/articles/5-1492). 16S rRNA FASTQ reads were trimmed at 220 nucleotides (nt) and 150 nt respectively with allowed expected errors of 2. ITS FASTQ reads were trimmed at 230 nt with an expected error of 2 and 4, respectively.

16S rRNA ASV sequences were subsequently classified using DADA2 against the SILVA database SSU Ref NR 99 release 138.1 (https://doi.org/10.5281/zenodo.4587955, https://www.ncbi.nlm.nih.gov/pubmed/23193283), and ITS ASV sequences were classified against the UNITE all eukaryotes general FASTA release version 8.2 [54] using a confidence threshold of 0.5.

All data analyses were performed in R (v.4.4.0) [55]. We used packages TreeSummarizedExperiment (v.2.12.0) [56], phyloseq (v.1.48.0) [57], and vegan (v.2.6-6.1) [58]. We generated plots using the ggplot2 package (v.3.5.1) [59] and edited them in Inkscape (v.0.92.4) [60]. We analysed samples that yielded at least 3000 reads (16S rRNA genes: n = 172, ITS2 sequences: n = 179, see Table S1 and S2).

To compare the number of bacterial and fungal ASVs detected in the model particle fractions and in soil, we randomly rarefied samples to 3015 (16S rRNA genes) and 5676 (ITS2 sequences) reads to establish equal sequencing depth across samples using the rrarefy function from the vegan package [58]. We performed differential abundance analysis using the ANCOMBC package (v.2.6.0) [61]. The analysis of microbiome compositions with bias correction determined which genera were significantly enriched and depleted in microbial communities associated with model particles compared to the “surrounding soil”, in which particles had been incubated. We examined temporal dynamics of particle-associated microbial communities over the incubation period with principal coordinate analysis (PCoA). Plots depict Aitchison distances, which are Euclidean distances calculated from centred-log-ratio transformed counts matrices and account for the compositional nature of amplicon sequencing data [62]. Additionally, we performed permutational analysis of variance (PERMANOVA) to test the significance of sampling time points on community composition using the adonis2 function of the vegan package.

## Results

### Microbial biomass increased on model particles over time, but not in the surrounding soil

We assessed microbial colonization and biomass of model particles using ddPCR and microscopic evaluation. Bacterial biomass increased on model particles over time as evidenced by increasing gene copy numbers on model chitin and cellulose particles over 50 and 47 days of incubation for lab and field incubations, respectively (Fig. 2A). After 50 days, the median number of 16S rRNA gene copies per gram of particles was highest on model chitin particles compared to cellulose and “no substrate” particles (Fig. 2A). Bacterial gene copies increased 52- and 31-fold on cellulose particles and 7- and 6-fold on chitin particles between day one and the end of the incubation. Microbial colonisation was lower for “no substrate” particles than chitin and cellulose particles, with gene copies increasing 3.6- and 4.5-fold more on chitin particles and 2.8-fold and 2.7-fold more on cellulose compared to “no substrate” particles after 50 and 47 days of incubation.

**Fig. 2.**
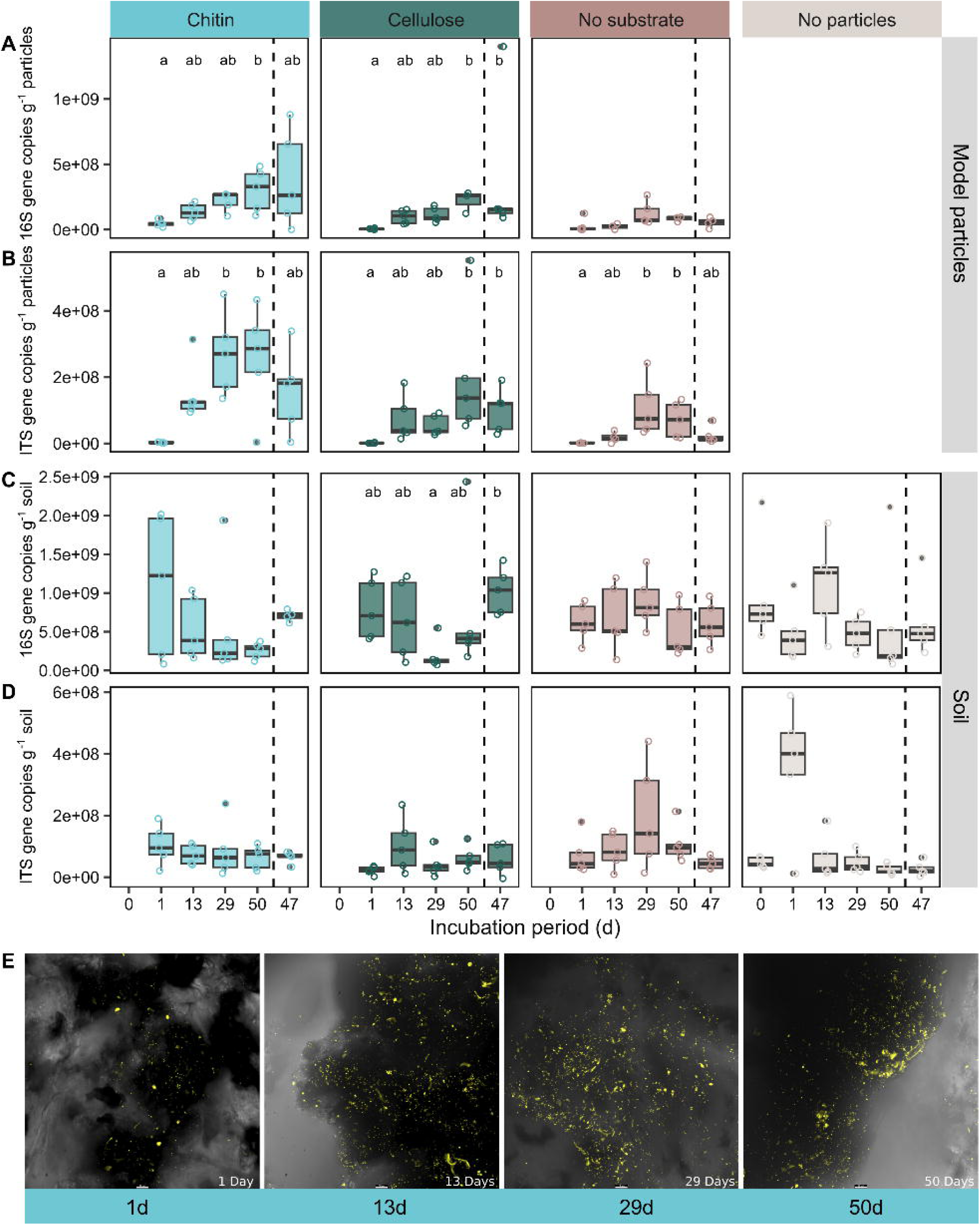
Bacterial, archaeal, and fungal biomass increased on model particles over the incubation period, but not in the surrounding soil. Boxplots show (**A, C**) 16S rRNA (16S) and (**B, D**) ITS gene copy numbers (per gram of particles or soil) (**A, B**) on model particles and (**C, D**) in the soil after zero, one, 13, 29, and 50 days of lab incubation, and 47 days of field incubation. Gene copy numbers can be interpreted as a proxy for microbial biomass associated with model particles or soil. (**E**) Fluorescence microscopy images show microbial DNA at model chitin particle surfaces after one, 13, 29, and 50 days of lab incubation. Yellow points represent DNA stained with SYBR Green I double stranded nucleic acid stain. Microscopy was performed using a Leica confocal laser scanning microscope coupled with a Thunder imaging system. Scale bars indicate 1 µm. (**A, B, C, D**) Different letters indicate significant pairwise differences across time points for each treatment and sample type based on Dunn’s post hoc tests with Bonferroni adjustment following Kruskal–Wallis tests (adjusted P-value < 0.05). Supplementary Table S3 gives the exact number of measurements that boxplots are based on.

The number of ITS gene copies was also the highest on chitin particles after 50 and 47 days of incubation (Fig. 2B). Between one and 50 days of lab incubation, we observed 143-fold, 152-fold, and 72-fold increases in ITS gene copies on chitin, cellulose, and “no substrate” particles (Fig. 2B). Fungal gene copies were 3.9- and 16.4-fold higher on chitin particles than on “no substrate” particles after 50 and 47 incubation days, and 1.9- and 10.9-fold higher on cellulose particles. Microscopy visualization of chitin particles supports the finding that particles were more densely populated after several days of incubation than after one day (Fig. 2E). Particles incubated in the lab and the field were similarly densely populated by microbes after 47 and 50 days. In contrast, microbial biomass in the soil surrounding the model particles, and biomass in the soil incubated without model particles did not reveal a clear pattern over time (Fig. 2C, 2D).

### ASV richness of model chitin particles decreased over the incubation period

To determine whether changes in microbial communities occurred on model particles and in soil across the different treatments, we investigated amplicon sequence variant (ASV) richness of rarefied data. Overall, the richness of initial particle-associated microbial communities was higher compared to communities associated with particles later in the incubation (Fig. 3). The number of bacterial taxa associated with model chitin particles dropped 1.2-fold between the start of the incubation and 50 and 47 days (Fig. 3A). Fungal richness of chitin particles was also approximately 1.2- and 1.5-fold lower after 50 and 47 days than after one day. In contrast, bacterial and fungal richness remained relatively unchanged on cellulose and “no substrate” particles between one and 50 days of incubation. Microbial richness of the incubated soil remained relatively constant across time points and treatments (Fig. 3).

**Fig. 3.**
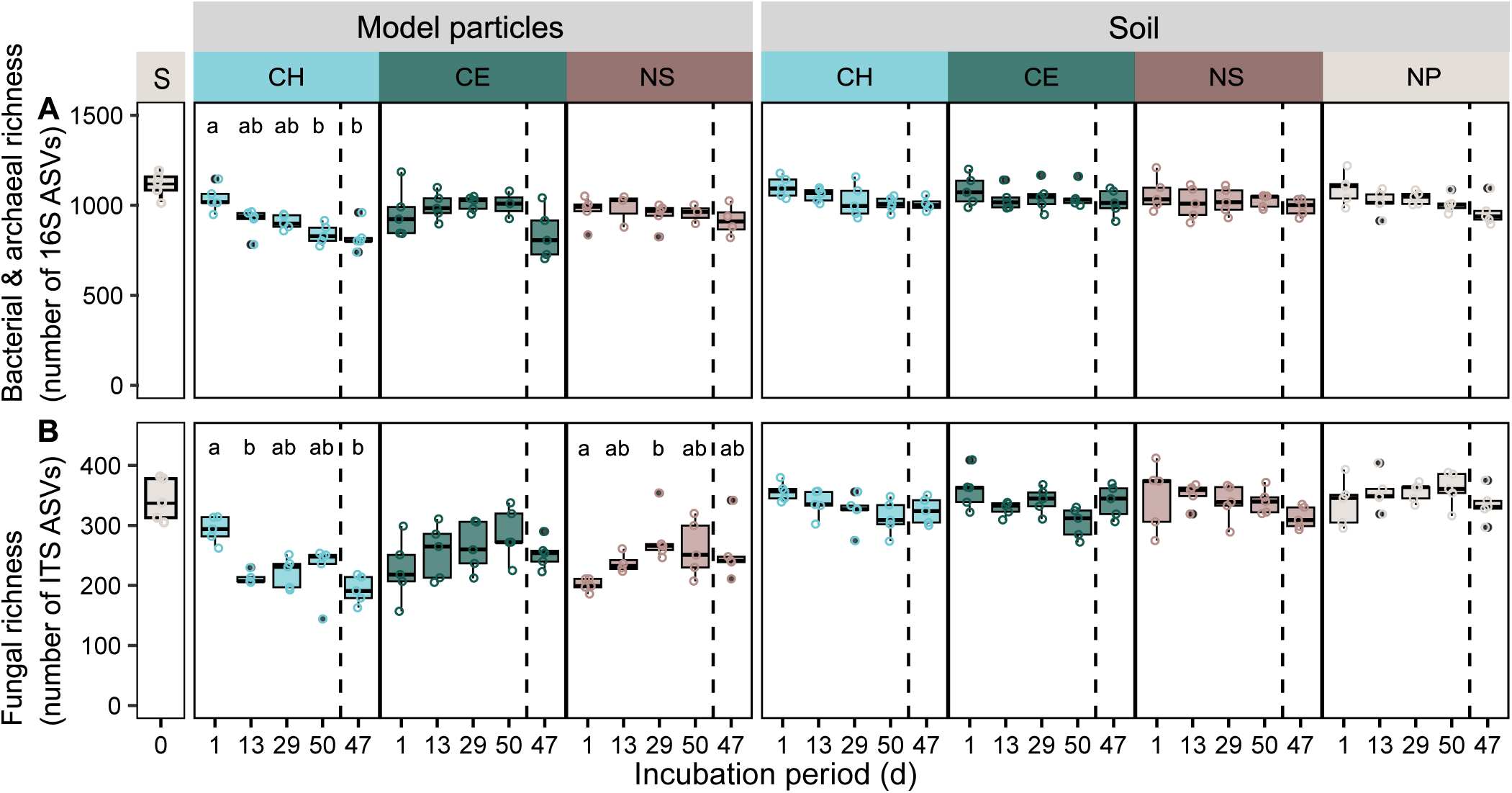
Bacterial and archaeal, and fungal richness of model chitin particles significantly decreased over the incubation period. Boxplots (from left to right) show (**A**) bacterial and archaeal richness (number of ASVs), and (**B**) fungal richness that was detected in (i) the soil, which was used to set up incubations, (ii) on model particles, and (iii) in the surrounding soil after one, 13, 29, and 50 days of lab incubation and 47 days field incubation. Sequencing data was randomly rarefied at the minimum read count for bacteria and archaea (3015 reads), and for fungi (5676 reads) to guarantee equal sequencing depth of samples. Different letters indicate significant pairwise differences across time points for each treatment and sample type based on Dunn’s post hoc tests with Bonferroni adjustment following Kruskal–Wallis tests (adjusted P-value < 0.05). S – sieved soil that was used to set up incubations, CH – chitin, CE – cellulose, NS – no substrate (control to account for the effect of the scaffold material of model particles, gellan gum), NP – soil without particles.

### Chitin particle-attached microbial communities underwent temporal changes

We investigated whether microbial community structure on model particles and the surrounding soil changed over the incubation period (Fig. 4). Differences in bacterial and fungal community composition between the time points were most pronounced for communities associated with chitin particles (Fig. 4A, 4D, Table S3). Communities detected on the first day of incubation were most similar to the communities of the soil used to set up incubations, while later communities (after 13, 29, and 50 days) differed more strongly from initial communities. PCoA plots suggest that bacterial and fungal communities associated with chitin particles, which had been incubated for 47 or 50 days, were very similar in their composition (Fig. 4A, 4D). In contrast, microbial communities associated with cellulose particles did not show pronounced changes over the course of the incubation (Fig. 4B, 4E). Bacterial communities associated with “no substrate” particles exhibited a shift in community composition after one day of incubation but remained similar over the remaining incubation period (Fig. 4C). In comparison, fungal communities associated with “no substrate” particles did not show a separation between the initial soil- and particle-associated communities after one incubation day compared to particles incubated for multiple days (Fig. 4F). Over all treatments, communities in soil did not exhibit strong changes over the incubation period (Fig. 4G-4L). Results of PERMANOVA analyses support the observation that sampling time point had a significant effect on composition of communities associated with chitin particles (Table S4).

**Fig. 4.**
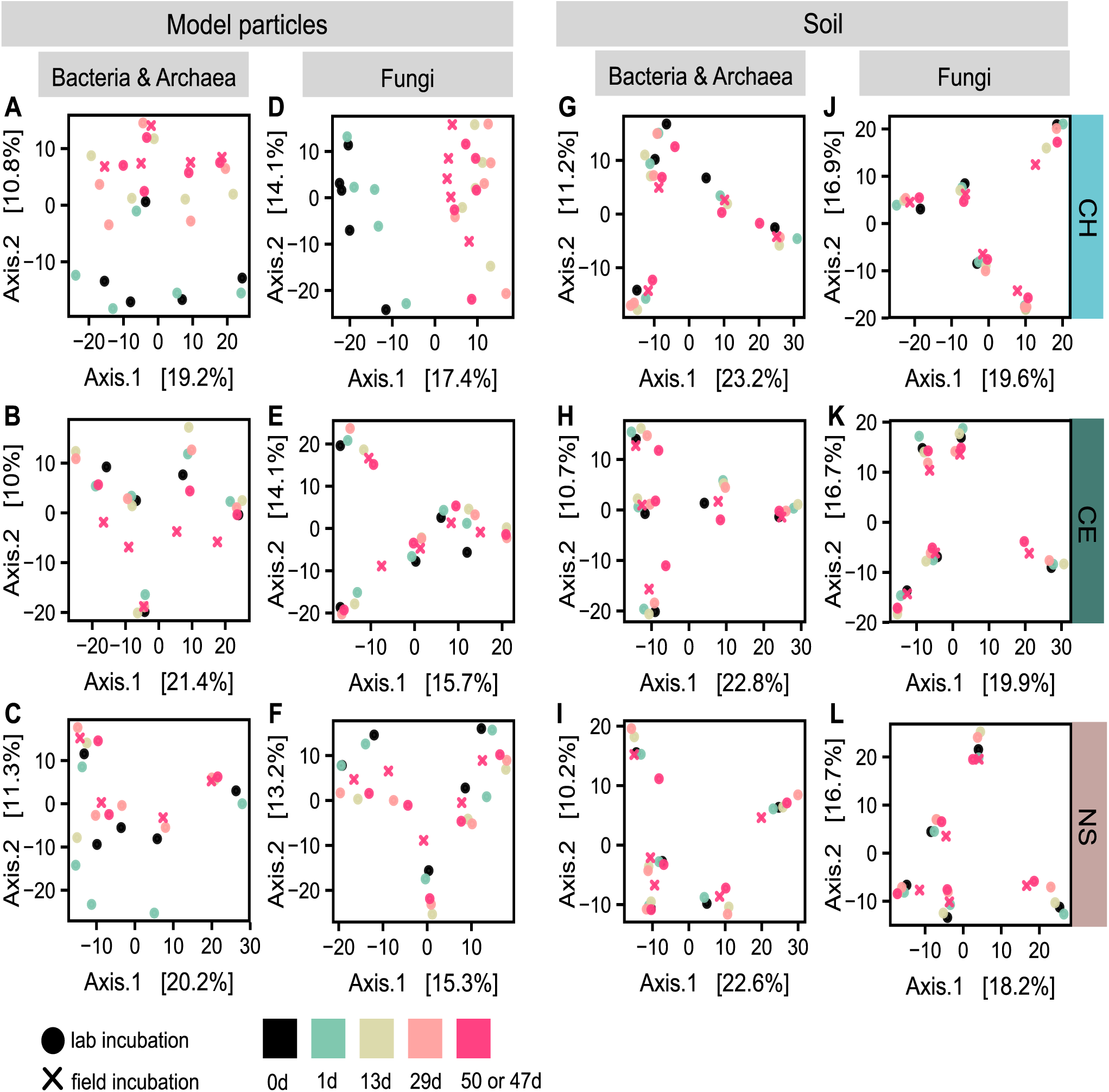
Chitin particle-associated bacterial and archaeal communities underwent temporal changes. PCoA plots show potential dissimilarity of (**A-C, G-I**) bacterial and archaeal, and (**D-F, J-L**) fungal communities (i) associated with model chitin, cellulose or “no substrate” particles or (ii) in the surrounding soil across harvest time points (day one to 50 or 47). Plots also include communities of the five sieved soils (at incubation start, 0d) that were used to set up mesocosms and mesh bags. PCoA plots are based on Aitchison distances (Euclidean distance of clr-transformed sequencing data). The colour of data points indicates the harvest time point, the shape of points depicts the type of incubation (lab or field). CH – chitin, CE – cellulose, NS – “no substrate”.

### Specific microbial genera were enriched on chitin and cellulose particles

Analysis of the initial soil community composition revealed bacterial genera *Mucilaginibacter*, *Acidothermus*, and *Bryobacter* and fungal genera *Mortierella*, *Russula*, and *Tomentella* among the most abundant genera, which served as the “seed bank” for microbial communities that could colonize model particles (Figure S1, S2). We compared relative abundances of microbial genera between model substrate particles and the surrounding soil to address whether substrates led to the enrichment of microbial genera. Differential abundance analysis showed enrichment of bacterial and fungal genera on chitin and cellulose particles compared to soil in which these particles had been incubated (Fig. 5, 6). At the same time, other genera were significantly depleted on model chitin and cellulose particles compared to the surrounding soil (Fig. S3, S4, S6, S7-). In the initial phase of colonization, few bacterial genera, including *Paenibacillus* and *Bacillus* on chitin particles and KD3-10, *Paenibacillus*, *Clostridium sensu stricto* 1, and *Micropepsis* on cellulose, were significantly more abundant on particles compared to the surrounding soil (Fig. 5A, 5B).

**Fig. 5.**
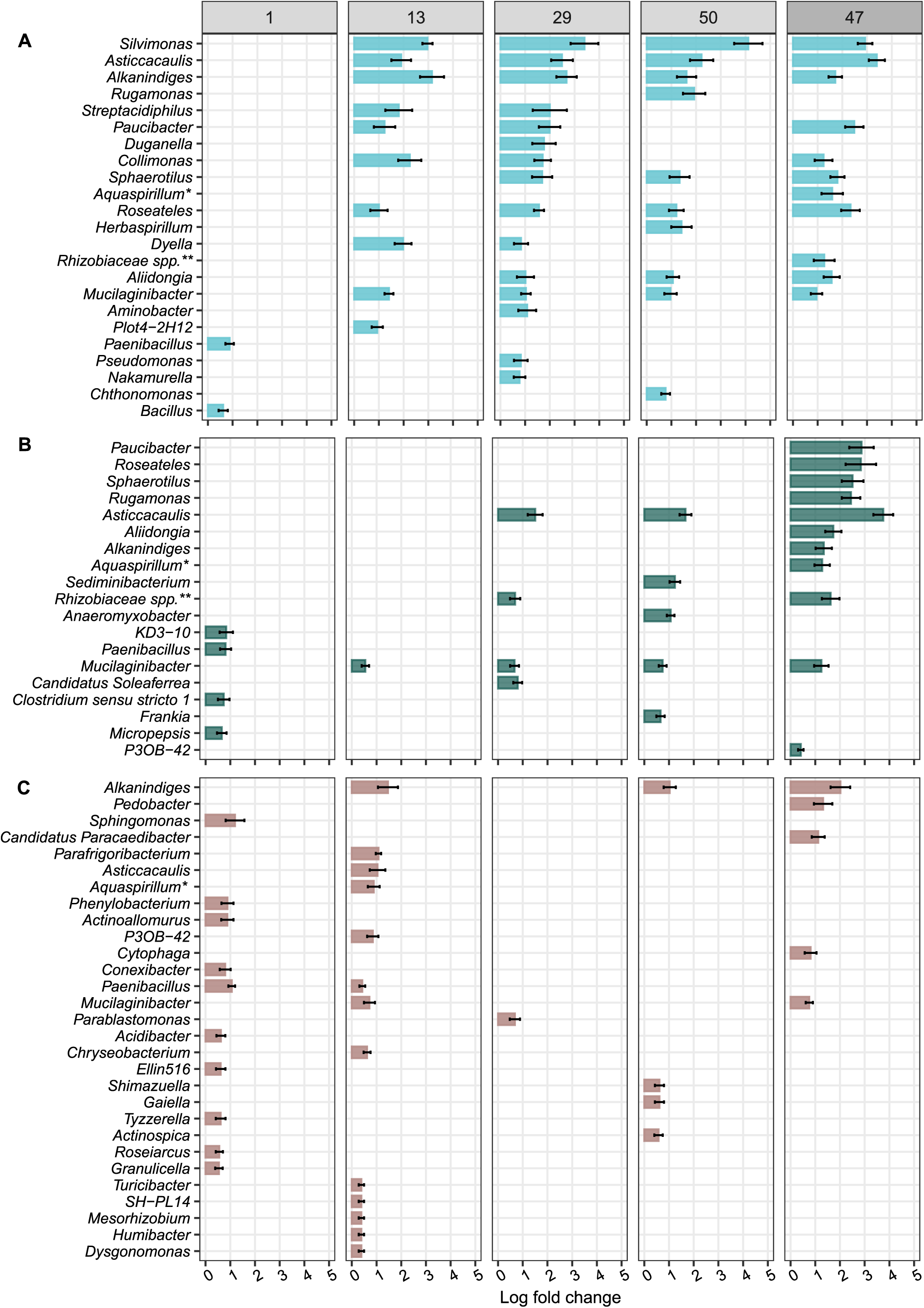
Model chitin and cellulose particles favoured certain bacterial genera compared to the soil, in which particles had been incubated. Plots show genera that were significantly enriched in their relative abundance on (**A**) model chitin, (**B**) model cellulose particles, and (**C**) “no substrate” model particles (controls) compared to the surrounding soil at one day, 13, 29, and 50 days after the start of the lab incubation and 47 days after the start of the field incubation. Bars represent the log fold change, a measure of effect size, in the relative abundance of significantly enriched genera (FDR-adjusted P value < 0.05). Differentially abundant genera were identified using the ANCOM-BC package [61]. Error bars represent the standard error of the log fold change. We controlled false discovery rates with the Benjamini-Hochberg multiple test correction. *Aquaspirillum\** refers to the *[Aquaspirillum] articum group*, *Rhizobiaceae spp.*\*\* refers to the *Allorhizobium-Neorhizobium-Pararhizobium-Rhizobium* clade. Note that we did not detect any genera that were significantly enriched on model “no substrate” particles after 47 days of incubation in the field. See Fig. S3-S5 for positive and negative log fold changes of genera on model particles compared to the surrounding soil.

**Fig. 6.**
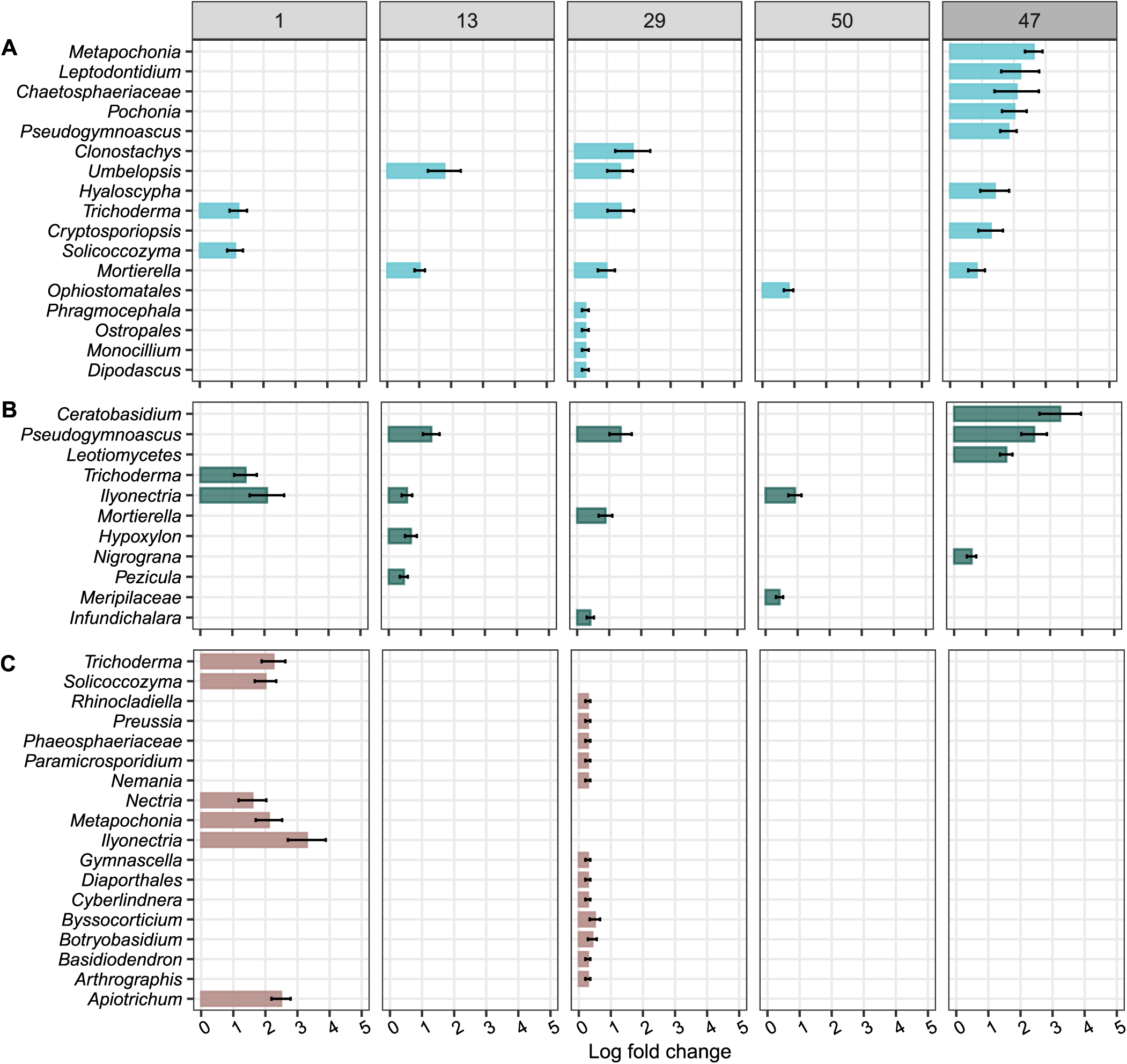
Model chitin and cellulose particles favoured certain fungal genera compared to the soil, in which particles had been incubated. Plots show genera which were significantly enriched in their relative abundance on (**A**) model chitin, (**B**) model cellulose particles, and (**C**) model “no substrate” particles (controls) compared to the surrounding soil at one day, 13, 29, and 50 days after the start of the lab incubation and 47 days after the start of the field incubation. Bars represent the log fold change, a measure of effect size, in the relative abundance of significantly enriched genera (FDR-adjusted P value < 0.05). Differentially abundant genera were identified using the ANCOM-BC package [61]. Error bars represent the standard error of the log fold change. We controlled false discovery rates with the Benjamini-Hochberg multiple test correction. Note that we did not detect any genera which were significantly enriched on model “no substrate” particles after 13 and 50 days of incubation in the lab, and 47 days of incubation in the field. See Fig. S6-S8 for positive and negative log fold changes of genera on model particles compared to the surrounding soil.

*Asticcacaulis* and *Mucilaginibacter* were two genera that were enriched on chitin particles at all later harvest time points (13d, 29d, 50d, 47d) (Fig. 5). Most genera were enriched sporadically at single time points. On chitin particles, bacterial genera *Alkanindiges*, *Silvimonas*, *Paucibacter*, *Roseatales*, and *Sphaerotilus* were amongst the most enriched at the later time points of the lab incubation (13d, 29d, 50d) (Fig. 5A). We show that a larger number of bacterial genera was significantly increased on chitin particles compared to soil than in the case of cellulose particles (Fig. 5). While the number of bacterial genera enriched on model chitin particles was similar for lab and field incubations after 47 and 50 days of incubation (Fig. 5A), more bacterial genera were enriched on cellulose particles incubated in the field than in the lab (Fig. 5B). Most bacterial genera enriched on chitin particles compared to soil were consistent in lab and field incubations (Fig. 5A, 5B).

We detected sporadic enrichment of fungal genera on particles compared to the surrounding soil (Fig. 6). Chitin particles were significantly enriched in *Umbelopsis* compared to surrounding soil (13d, 29d) (Fig. 6A). *Pseudogymnascus* was relatively more abundant on cellulose particles than in the surrounding soil after 13 and 47 days of incubation (Fig. 6B). We found a larger number of fungal genera enriched on chitin and cellulose particles incubated in the field (after 47 days) than in the lab (after 50 days) (Fig. 6A). For example, *Metapochonia*, *Pochonia*, and *Pseudogymnascus* were only significantly enriched on chitin particles, as well as *Ceratobisidium* and *Leotiomycetes* on cellulose particles in the field. Overall, fewer fungal than bacterial genera were significantly enriched on model chitin particles compared to soil (Fig. 6A, 5A).

## Discussion

Our study applied an innovative approach toward investigating microbial community assemblages directly at the physical interface of POM. We studied temporal changes in microbial communities colonizing model substrate particles under both lab and field conditions, revealing enrichment of specific taxa at the particle-scale over time. Our study demonstrates model substrate particles as a suitable tool for studying substrate-specific microbial communities on POM in soil. We demonstrate that microbial colonization and temporal change in microbial community structure on the surface of model particles are substrate-specific. The higher number of gene copies on substrate particles compared to “no substrate” particles for bacteria and fungi indicates that presence of substrates enhanced microbial colonization and growth on particles. Our results suggest that microbes were physically attached to particles, as they would otherwise have been removed during the washing steps. This is in line with a previous study which proposed that bacteria are widely surface-attached in soil to improve enzymatic return on investment and promote microbial growth [8].

Analysis of changes in ASV number revealed a decrease in bacterial and fungal richness on model chitin particles over time. In contrast, bacterial richness remained unchanged on cellulose and “no substrate” particles during the incubation period, while fungal richness increased on both model particle types. Microbial communities associated with chitin particles underwent temporal changes, whereas cellulose particles did not exhibit clear temporal patterns. Communities associated with chitin particles became more chitin-specific over time, supporting the occurrence of temporal succession on chitin-containing POM [63]. Cellulose did not select for specific microbial communities as strongly as chitin, potentially due to the greater chemical complexity of chitin, which may necessitate a specific suite of enzymes or microbial interactions for degradation. In contrast, we did not detect temporal patterns of microbial community composition for the soil incubated with chitin particles, suggesting that the effects of complex particulate substrates on microbial communities may not be captured with composite soil samples.

Fewer bacterial genera were enriched on model cellulose particles than on chitin particles in the lab incubation, supporting our claim that chitin exerts a stronger selective pressure on microbial communities than cellulose. Genera enriched on chitin and cellulose were potentially well-adapted to utilize or degrade chitin, cellulose or their degradation by-products. Members of some genera that were enriched on model substrate particles have been described to possess the genetic potential to produce chitin- or cellulose-degrading enzymes. For example, the genus *Asticcacaulis* contains many glycoside hydrolases, key enzymes targeting chitin [15], while members of *Mucilaginibacter* may possess certain glycoside hydrolases capable of degrading chitin and cellulose [64, 65]. Species of other significantly enriched genera on chitin and/or cellulose particles, *Silvimonas* and *Collimonas,* have been found to possess chitinases [66–68] and a member of *Roseateles,* which was enhanced on cellulose particles in the field, is known to degrade cellulose [69].

However, not all taxa detected on model substrate particles necessarily degraded the substrate themselves. Some taxa might have profited from degradation products released by other microbes or might have fed on metabolites of degraders. Importantly, degradation potential is not necessarily conserved at the genus level, and assigning specific taxa to substrate degradation is limited based on amplicon sequencing results. Some genera enriched on chitin and cellulose particles were also enriched on “no-substrate” particles (Fig. 5), indicating non-specific colonization or a broader enzymatic repertoire. For instance, *Mucilaginibacter* was enriched on chitin particles, cellulose particles, and “no substrate” particles (Fig. 5), suggesting that members of this genus were not directly favoured by chitin or cellulose but may have utilized other particle components, or stochastically colonized particle surfaces. Similarly, enrichment of *Paenibacillus* on all particle types after one incubation day suggests efficient colonization of particles irrespective of substrate, although the genus includes known chitin and cellulose degraders [70–72].

Comparison of chitin particles to “no substrate” particles (Fig. S9) supports our observation that bacterial genera like *Silvimonas, Asticcacaulis, Alkanindiges, Collimonas*, and *Mucilaginibacter* were favoured by the presence of chitin rather than particle surfaces or other particle components. To a lesser extent, enrichment of *Asticcacaulis* and *Collimonas* could be attributed to the presence of cellulose, as these were also significantly enhanced on cellulose particles compared to “no substrate” particles (Fig. S10). We recommend including “no substrate” particles as controls for particles containing specific substrates. Overall, it would be advantageous for future studies to investigate genes and extracellular enzyme activity of taxa associated with model substrate particles or perform isotopic tracing experiments in combination with model particles to uncover taxa that degrade specific substrates.

Our results demonstrate that changes in microbial communities are indeed more pronounced at the particle interface than at the soil scale. Only a small number of fungal genera, including *Umbelopsis* and *Pseudogymnascus*, significantly increased on model chitin and cellulose substrate particles compared to the surrounding soil. Both genera were significantly enriched on substrate particles compared to “no substrate” controls (Fig. S11, S12). This suggests that members of *Umbelopsis* may degrade chitin or utilize its degradation products, while *Pseudogymnascus* may similarly use cellulose-derived compounds. Nevertheless, our results support that the ability to degrade chitin or cellulose is potentially more widespread among fungal than bacterial taxa [12, 73]. Consequently, these specific substrates might exhibit a lower selective pressure on fungi than on bacteria. Comparisons between model substrate particles and ”no substrate” particles underline that differences in relative abundances of microbial genera were driven by the specific substrate and likely not other particle components (Fig. S9-S12). We hypothesized that microbial communities in lab and field incubations would follow similar community assembly trajectories driven by substrate type. This was partially supported, as a subset of taxa was shared between the lab and field experiments and consistently showed substrate-specific enrichment. However, enrichment patterns differed between lab and field. In the lab, more bacterial genera were enriched on chitin particles than on cellulose, whereas equal numbers were enriched on both substrates in the field. For fungi, substantially more genera were enriched on model chitin and cellulose particles in the field than the lab. For example, *Metapochonia* and *Pochonia* were both only significantly enriched on chitin particles in the field (Fig. 6A), suggesting either migration to field mesocosms during the 47-day incubation or enhanced growth under field conditions. For bacteria and fungi, log fold changes were larger for field-incubated particles than for lab-incubated particles. While mesocosms containing model particles and soil were incubated under constant temperature and moisture conditions, the contents of mesh bags in the field were exposed to natural fluctuations and potential influxes from plant exudates over the incubation period. Interpretation of substrate-specific enrichment is further constrained by differences in sampling design, as lab incubation captured multiple harvests while the field experiment represents a single point in time. These differences highlight the need for caution when translating lab findings to nature.

Our study underpins that model substrate particles provide a highly flexible, standardized experimental system for the investigation of microbial degrader communities [31, 38, 74]. Model particles allow control over particle size, shape, and substrate composition [38], and remain stable and retrievable through incubation unlike natural substrates that become indistinguishable from soil over time. Particles can also be investigated using microscopy due to the high transparency. Future experiments could be conducted using particles to isolate substrate-specific consortia for downstream microbiological applications.

A potential caveat of this method is the use of gellan gum, which may be metabolized by a small subset of microorganisms, despite its resistance to enzymatic degradation [36, 75]. The low increase in gene copies on “no substrate” particles indicates non-specific colonization. Only few genera including *Paenibacillus*, *Alkanindiges, Trichoderma,* and *Ilyonectria* were enriched across all particle types, whereas most genera enriched on model chitin and cellulose particles were not enriched on “no substrate” particle controls. It is essential to bear in mind that a certain fraction of model particle-associated microbes was possibly lost due to the washing steps or that passively attached communities were captured during sequencing. It might be advisable to shorten the here applied incubation duration in future experiments to capture and study more fine-scale temporal dynamics of microbial communities associated with POM in soil.

## Conclusion

We found that complex substrates, chitin and cellulose, selected for specific microbial communities. Microbial communities associated with chitin underwent stronger temporal changes than those associated with cellulose, demonstrating substrate-specificity of communities in the here investigated forest soil. This approach has the potential to advance our knowledge of microbial taxa involved in degradation of complex biopolymers in soil by enabling isolation and characterization of substrate-specific microbial communities. We emphasize that further studies are necessary to determine whether these patterns are generalisable to other soil types and ecosystems. This study demonstrates the applicability of substrate particles as a model system for investigating substrate-specific microbial communities associated with particulate organic matter and their temporal dynamics in soil.

## Acknowledgements

This research was funded by the European Research Council under the European Union’s Horizon 2020 research and innovation program (grant agreement No. 819446). Lauren Alteio is funded by the European Union’s Horizon 2020 research and innovation programme under grant agreement No. 101060698. The authors would like to thank Benjamin Zwirzitz for his helpful advice and discussions throughout the course of this work. Marc Mussmann was supported by the Austrian Science Fund (FWF) project number P31010. We thank Sean Darcy, Ksenia Guseva, and Christian Ranits for their help in the field, and Petra Pjevac for advising and coordinating DNA extraction and amplicon sequencing. We additionally acknowledge the Life Science Computer Cluster (LiSC) of the University of Vienna for enabling the processing of the raw amplicon sequencing data.

## Credit authorship contribution statement

**Eva Simon**: Formal analysis, Investigation, Visualization, Writing – original draft, Writing – review & editing. **Lauren Alteio**: Conceptualization, Formal analysis, Investigation, Methodology, Project administration, Writing – original draft, Writing – review & editing. **Alexander König:** Investigation, Writing – review & editing. **Bruna Imai**: Investigation, Writing – review & editing. **Julia Horak:** Investigation, Writing – review & editing. **Julia Wiesenbauer**: Investigation, Writing – review & editing. **Joana Séneca**: Data curation, Writing – review & editing. **Bela Hausmann**: Data curation, Writing – review & editing. **Marc Mussmann**: Methodology, Writing – review & editing. **Barbara Kitzler:** Resources, Writing – review & editing. **Christina Kaiser**: Conceptualization, Funding acquisition, Project administration, Supervision, Writing – review & editing.

## Data Availability

Raw 16S rRNA genes and ITS2 amplicon sequencing data have been deposited in the NCBI Sequence Read Archive under the BioProject accession number PRJNA1208268. Bacterial, archaeal, and fungal gene copy numbers are available on the Zenodo repository (10.5281/zenodo.17901929).

