## Supplemental File for "Model substrate particles uncover dynamics of microbial communities associated with particulate organic matter decomposition in soil"

Supplementary Material

Table S1 Number of samples that were analysed in 16S rRNA gene sequencing analysis after excluding all samples with less than 3000 sequencing reads. CH – chitin, CE – cellulose, NS – “no substrate”, NP – no particle soil.

| Incubation period (d) | Model particles | | | Soil | | | |
| --- | --- | --- | --- | --- | --- | --- | --- |
|  | CH | CE | NS | CH | CE | NS | NP |
| 0 | - | - | - | - | - | - | 5 |
| 1 | 5 | 5 | 5 | 5 | 5 | 5 | 5 |
| 13 | 5 | 5 | 3 | 5 | 5 | 5 | 5 |
| 29 | 5 | 5 | 5 | 5 | 5 | 5 | 5 |
| 50 | 5 | 3 | 3 | 5 | 5 | 5 | 5 |
| 47 | 5 | 5 | 4 | 4 | 5 | 5 | 5 |

Table S2 Number of samples that were analysed in ITS2 sequences sequencing analysis after excluding all samples with less than 3000 sequencing reads. CH – chitin, CE – cellulose, NS – “no substrate”, NP – no particle soil.

| Incubation period (d) | Model particles | | | Soil | | | |
| --- | --- | --- | --- | --- | --- | --- | --- |
|  | CH | CE | NS | CH | CE | NS | NP |
| 0 | - | - | - | - | - | - | 5 |
| 1 | 5 | 5 | 5 | 5 | 5 | 5 | 5 |
| 13 | 5 | 5 | 4 | 5 | 5 | 5 | 5 |
| 29 | 5 | 5 | 5 | 5 | 5 | 5 | 5 |
| 50 | 5 | 5 | 5 | 5 | 5 | 5 | 5 |
| 47 | 5 | 5 | 5 | 5 | 5 | 5 | 5 |

Table S3 Number of measurements that underlie boxes in Fig. 2. CH – chitin, CE – cellulose, NS – “no substrate”, NP – no particle soil.

|  | Incubation period (d) | Model particles | | | Soil | | | |
| --- | --- | --- | --- | --- | --- | --- | --- | --- |
|  |  | CH | CE | NS | CH | CE | NS | NP |
| 16S | 0 | - | - | - | - | - | - | 5 |
|  | 1 | 5 | 5 | 5 | 5 | 5 | 5 | 5 |
|  | 13 | 5 | 5 | 3 | 5 | 5 | 5 | 5 |
|  | 29 | 5 | 5 | 5 | 5 | 5 | 5 | 5 |
|  | 50 | 5 | 3 | 3 | 5 | 5 | 5 | 5 |
|  | 47 | 5 | 5 | 4 | 4 | 5 | 5 | 5 |
| ITS2 | 0 | - | - | - | - | - | - | 5 |
|  | 1 | 5 | 5 | 5 | 5 | 5 | 5 | 5 |
|  | 13 | 5 | 5 | 4 | 5 | 5 | 5 | 5 |
|  | 29 | 5 | 5 | 5 | 5 | 5 | 5 | 5 |
|  | 50 | 5 | 5 | 5 | 5 | 5 | 5 | 5 |
|  | 47 | 5 | 5 | 5 | 5 | 5 | 5 | 5 |

Table S4 Results of PERMANOVA of Euclidean distances testing the effect of incubation period on bacterial and archaeal (16S rRNA) and fungal (ITS2) community composition within sample types. Analyses were performed with the adonis2 function (vegan package) with 9999 permutations on Euclidean distances of clr-transformed composition data. CH – chitin, CE – cellulose, NS – “no substrate”; Df – degrees of freedom, SS – sum of squares, F – F statistic, P-value. Significance levels: *** P < 0.001, ** P < 0.01, * P < 0.05, . P = 0.05.

| Treatment | Type | df | SS | R^2^ | F | P-value | Significance |
| --- | --- | --- | --- | --- | --- | --- | --- |
| 16S rRNA | | | | | | | |
| CH | **particles** | **4** | **5261.5** | **0.18** | **1.40** | **0.01** | ***** |
|  | soil | 4 | 3438.3 | 0.11 | 0.81 | 0.87 |  |
| CE | particles | 4 | 4427 | 0.14 | 0.99 | 0.45 |  |
|  | soil | 4 | 3378.1 | 0.11 | 0.78 | 0.92 |  |
| NS | particles | 4 | 4442.2 | 0.16 | 1.01 | 0.41 |  |
|  | soil | 4 | 3195.1 | 0.10 | 0.76 | 0.95 |  |
| ITS2 | | | | | | | |
| CH | **particles** | **4** | **6768.9** | **0.24** | **1.99** | **0.0003** | ******* |
|  | soil | 4 | 3505.7 | 0.12 | 0.88 | 0.69 |  |
| CE | **particles** | **4** | **5183.5** | **0.17** | **1.29** | **0.05** | **.** |
|  | soil | 4 | 3462.8 | 0.11 | 0.84 | 0.76 |  |
| NS | **particles** | **4** | **5013.7** | **0.17** | **1.29** | **0.05** | **.** |
|  | soil | 4 | 3554.1 | 0.12 | 0.92 | 0.61 |  |

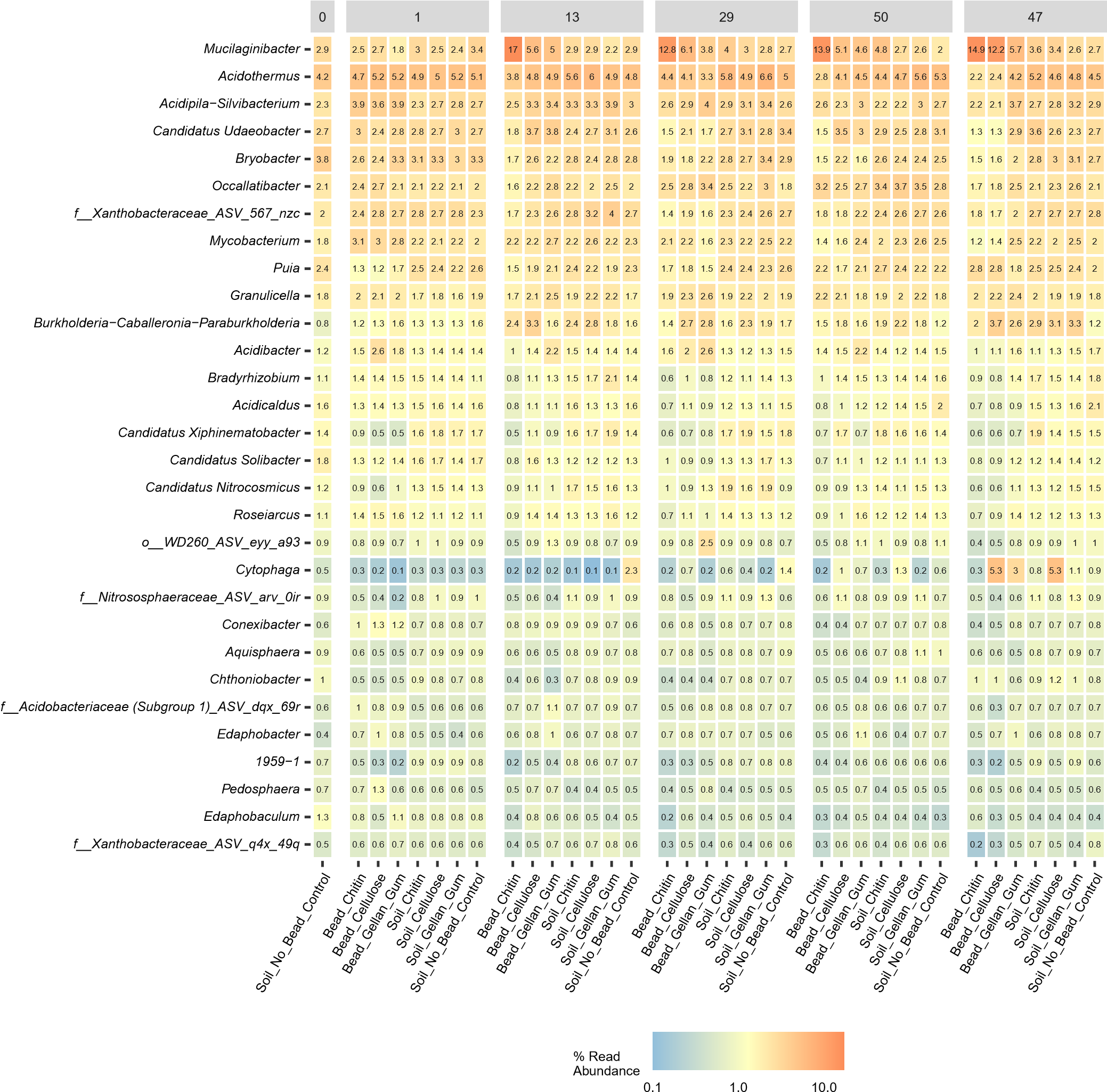

Fig. S1 Genus-level heatmaps show the relative abundance of the 30 most abundant bacterial genera across treatments and sampling days. Taxonomic profiles were aggregates at the genus level using ampvis2. Samples are grouped by treatment and faceted by days from the start of the experiment. Colour intensity represents log_10_-transformed relative abundances. Individual relative abundances values are printed on cells.

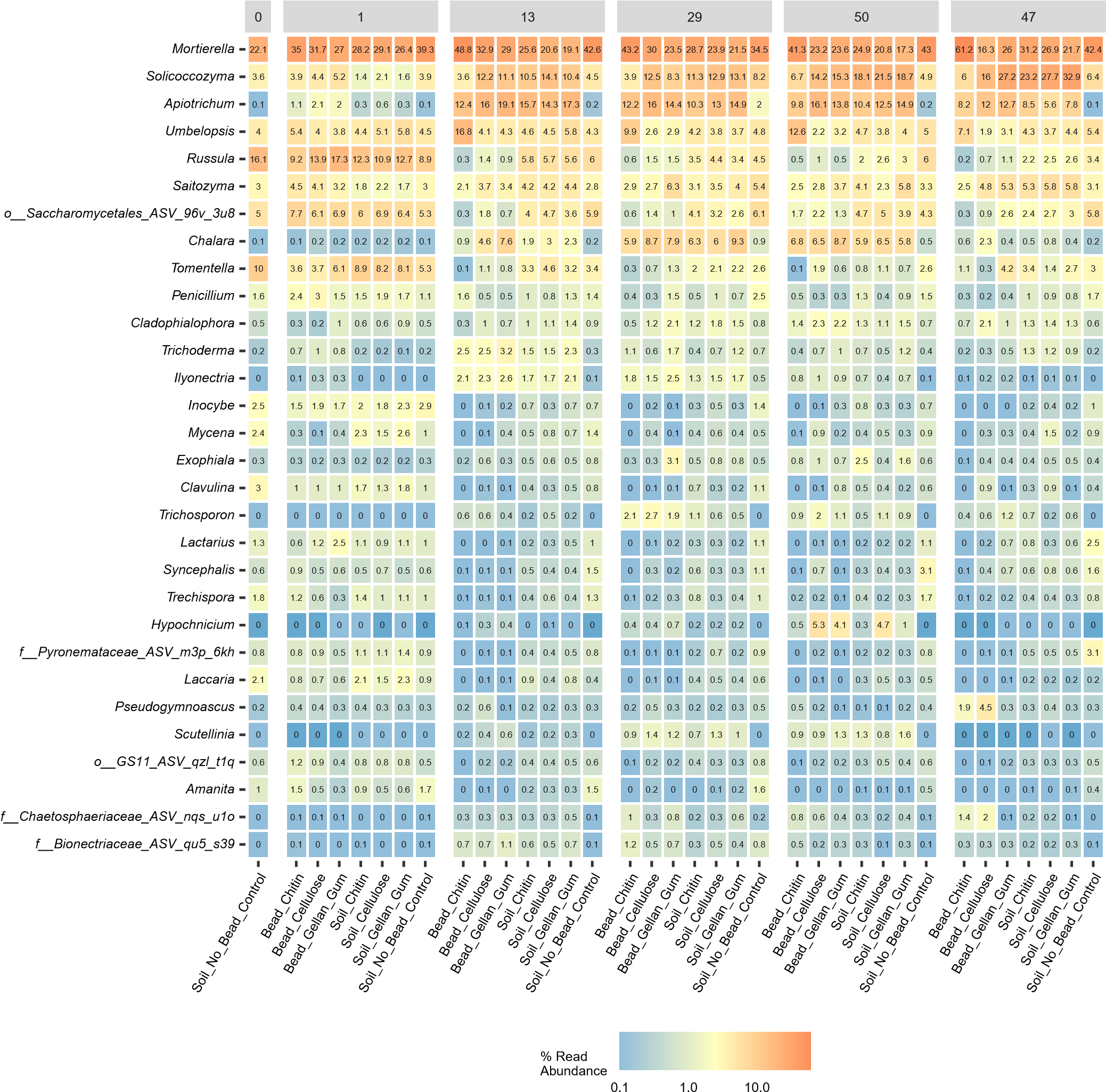

Fig. S2 Genus-level heatmaps show the relative abundance of the 30 most abundant fungal genera across treatments and sampling days. Taxonomic profiles were aggregates at the genus level using ampvis2. Samples are grouped by treatment and faceted by days from the start of the experiment. Colour intensity represents log_10_-transformed relative abundances. Individual relative abundances values are printed on cells.

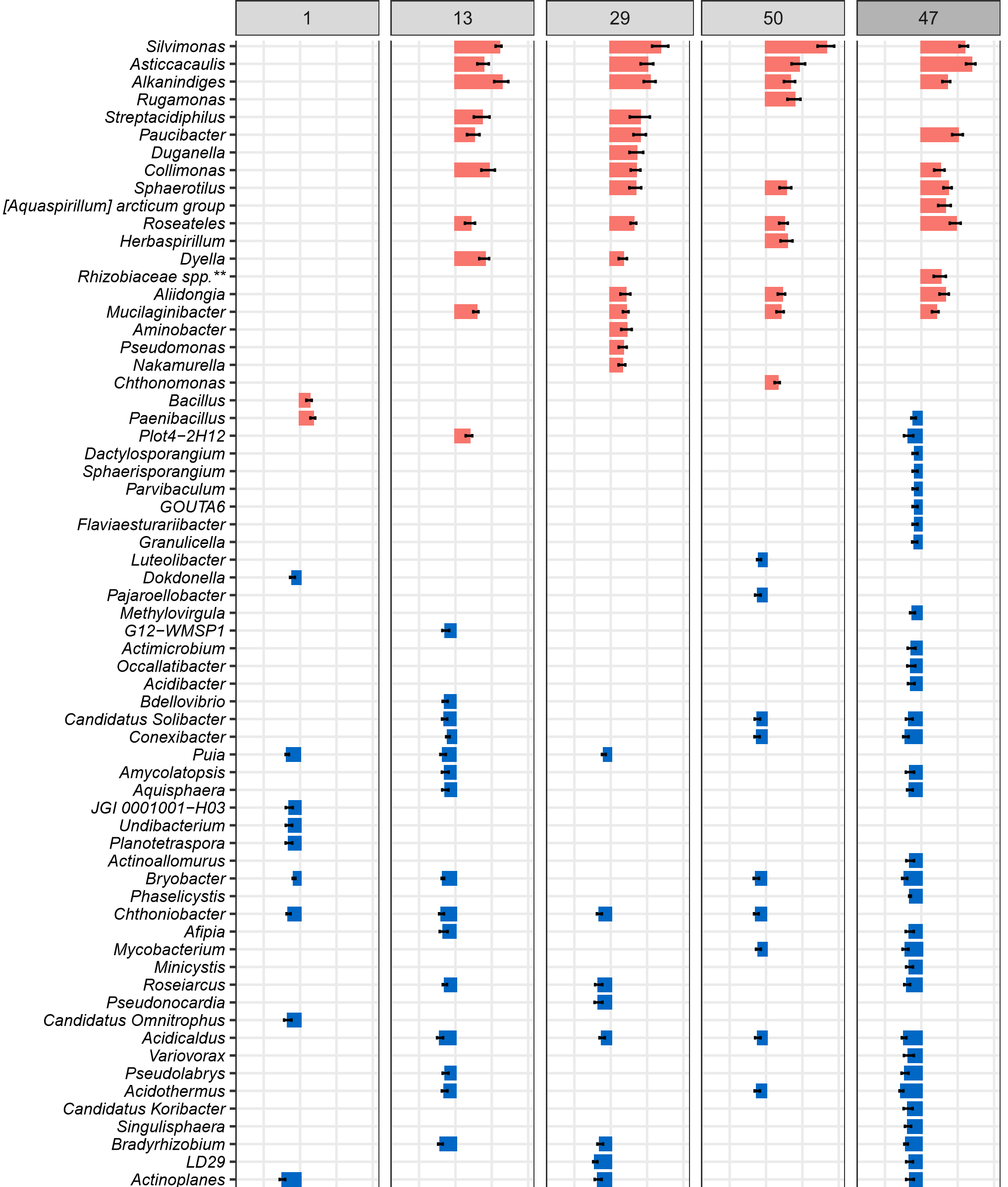

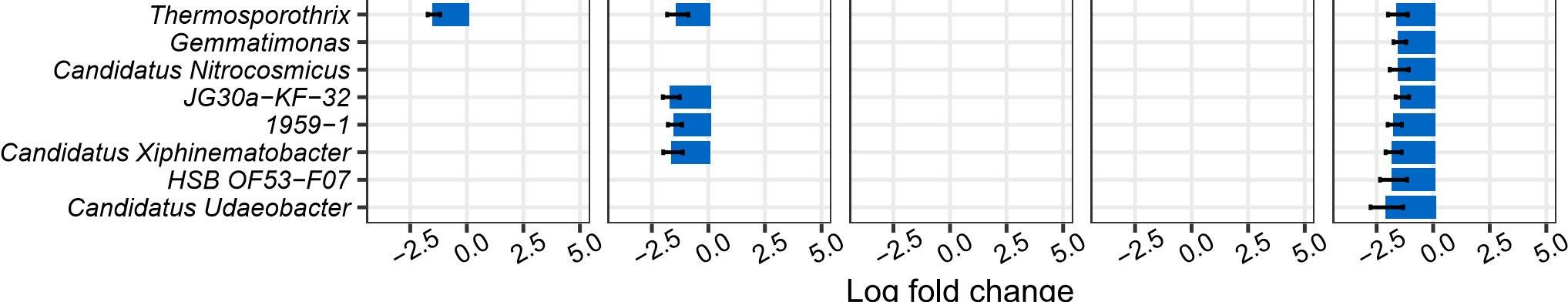
Fig. S3 Plots show bacterial and archaeal genera which were significantly enriched (positive log fold changes) and significantly depleted (negative log fold changes) on model chitin particles compared to the soil, in which particles had been incubated, one, 13, 29, and 50 days after the start of the lab incubation and 47 days after the start of the field incubation. Log fold changes are calculated from relative abundance data. Bars represent the positive or negative log fold change, a measure of effect size, in the relative abundance of significantly enriched genera (FDR-adjusted P value < 0.05). Differentially abundant genera were identified using the ANCOM-BC package (Lin and Peddada, 2020). Error bars represent the standard error of the log fold change. We controlled for false discovery rates with the Benjamini-Hochberg multiple test correction. *Rhizobiaceae spp.*** refers to the *Allorhizobium-Neorhizobium-Pararhizobium-Rhizobium* clade.

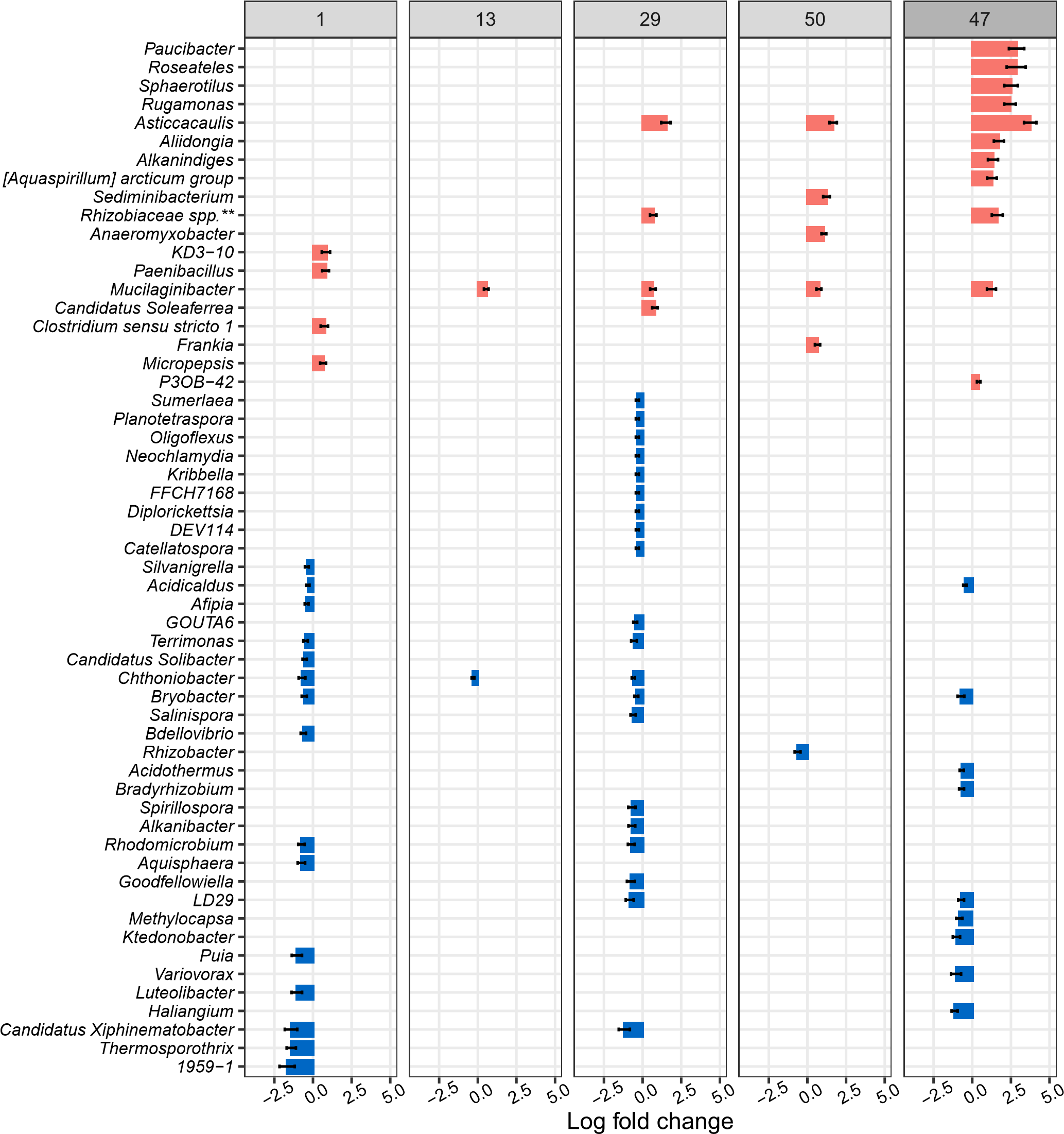
Fig. S4 Plots show bacterial and archaeal genera which were significantly enriched (positive log fold changes) and significantly depleted (negative log fold changes) on model cellulose particles compared to soil, in which particles had been incubated, one, 13, 29, and 50 days after the start of the lab incubation and 47 days after the start of the field incubation. Log fold changes are calculated from relative abundance data. Bars represent the positive or negative log fold change, a measure of effect size, in the relative abundance of significantly enriched genera (FDR-adjusted P-value < 0.05). Differentially abundant genera were identified using the ANCOM-BC package (Lin and Peddada, 2020). Error bars represent the standard error of the log fold change. We controlled for false discovery rates with the Benjamini-Hochberg multiple test correction. *Rhizobiaceae spp.*** refers to the *Allorhizobium-Neorhizobium-Pararhizobium-Rhizobium* clade.

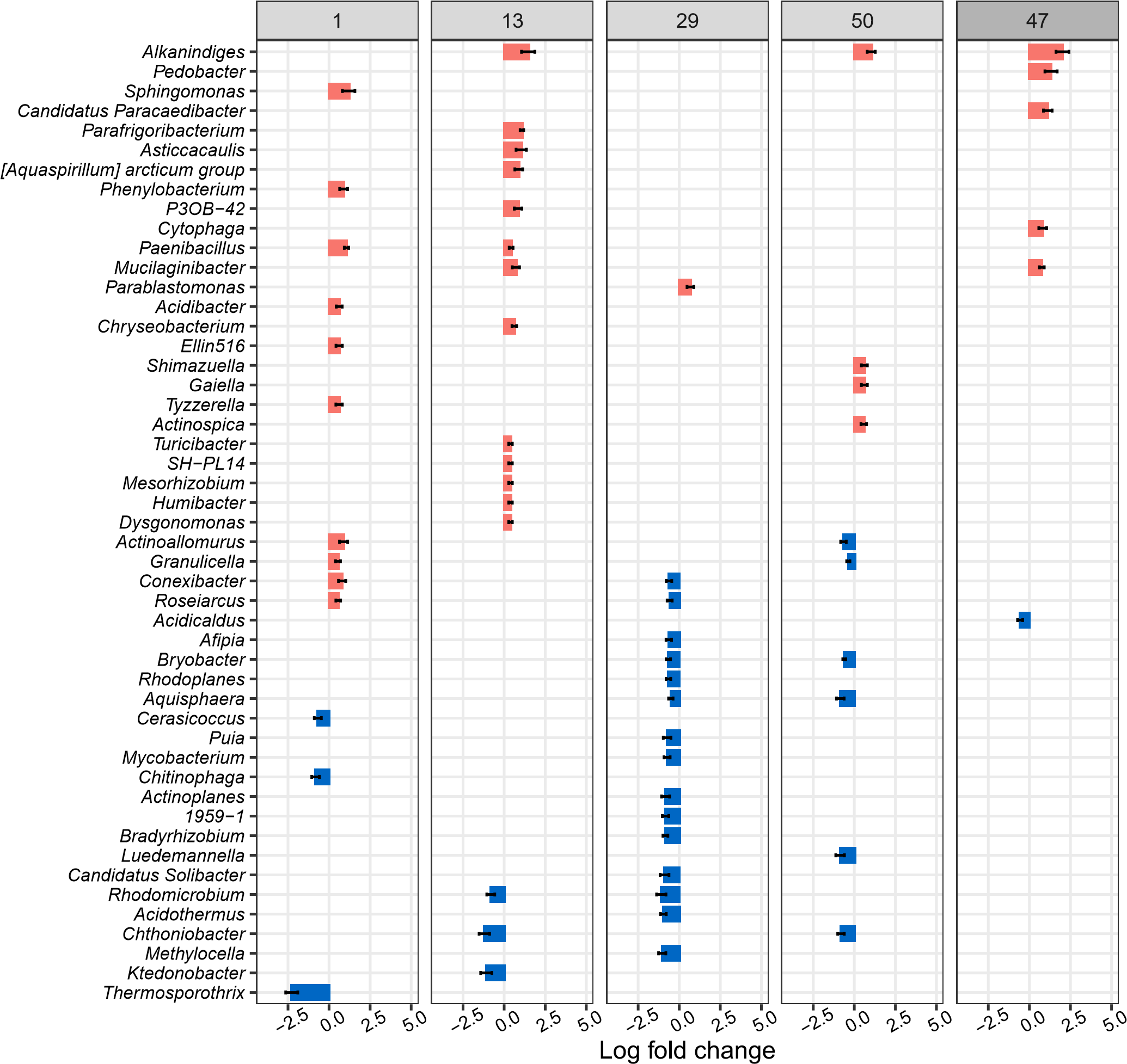

Fig. S5 Plots show bacterial and archaeal genera that were significantly enriched (positive log fold changes) and significantly depleted (negative log fold changes) on “no substrate” model particles compared to the soil, in which particles had been incubated, one, 13, 29, and 50 days after the start of the lab incubation and 47 days after the start of the field incubation. Log fold changes are calculated from relative abundance data. Bars represent the positive or negative log fold change, a measure of effect size, in the relative abundance of significantly enriched genera (FDR-adjusted P-value < 0.05). Differentially abundant genera were identified using the ANCOM-BC package (Lin and Peddada, 2020). Error bars represent the standard error of the log fold change. We controlled for false discovery rates with the Benjamini-Hochberg multiple test correction.

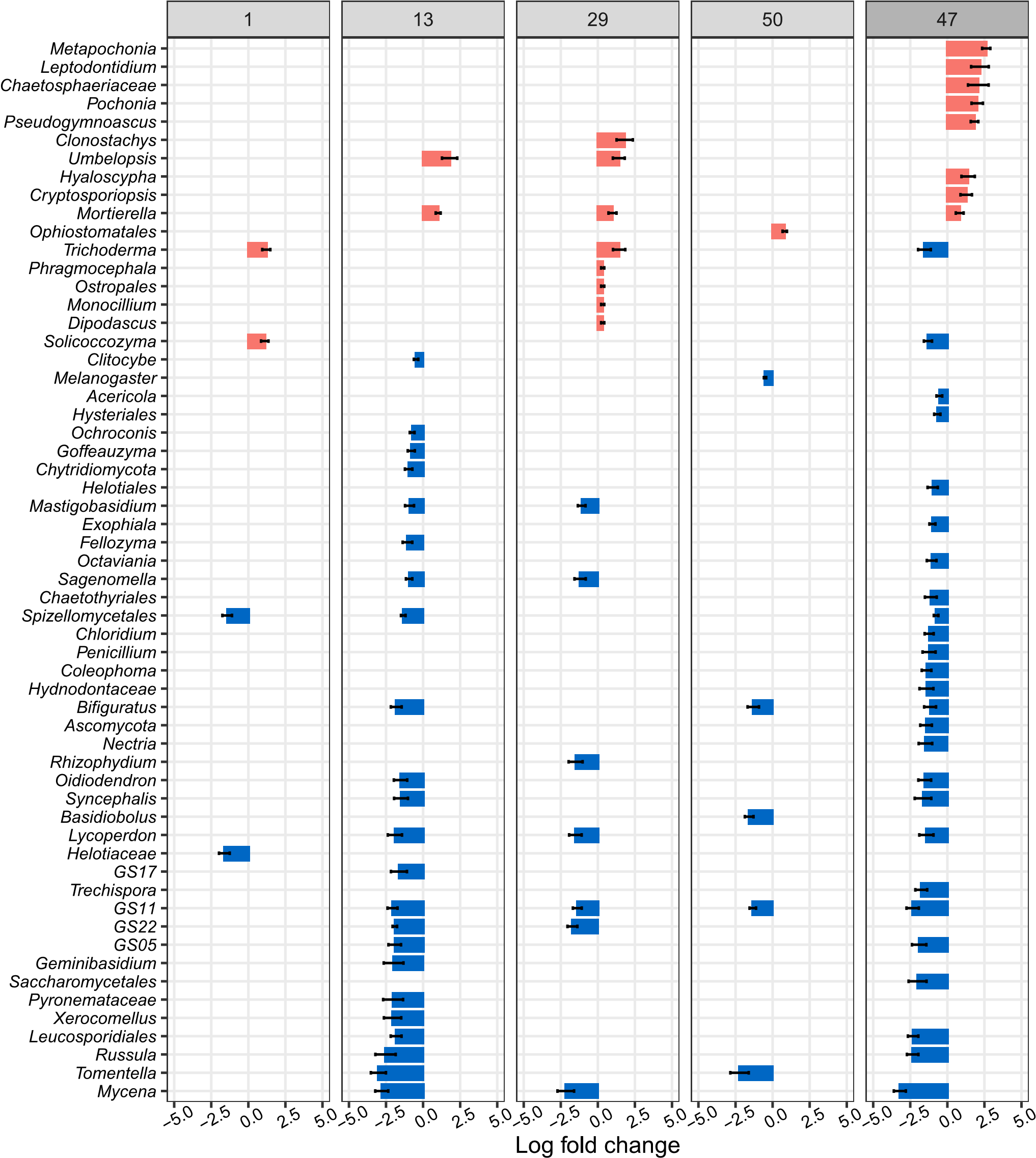

Fig. S6 Plots show fungal genera which were significantly enriched (positive log fold changes) and significantly depleted (negative log fold changes) on model chitin particles compared to the soil, in which particles had been incubated, one, 13, 29, and 50 days after the start of the lab incubation and 47 days after the start of the field incubation. Log fold changes are calculated from relative abundance data. Bars represent the positive or negative log fold change, a measure of effect size, in the relative abundance of significantly enriched genera (FDR-adjusted P-value < 0.05). Differentially abundant genera were identified using the ANCOM-BC package (Lin and Peddada, 2020). Error bars represent the standard error of the log fold change. We controlled for false discovery rates with the Benjamini-Hochberg multiple test correction.

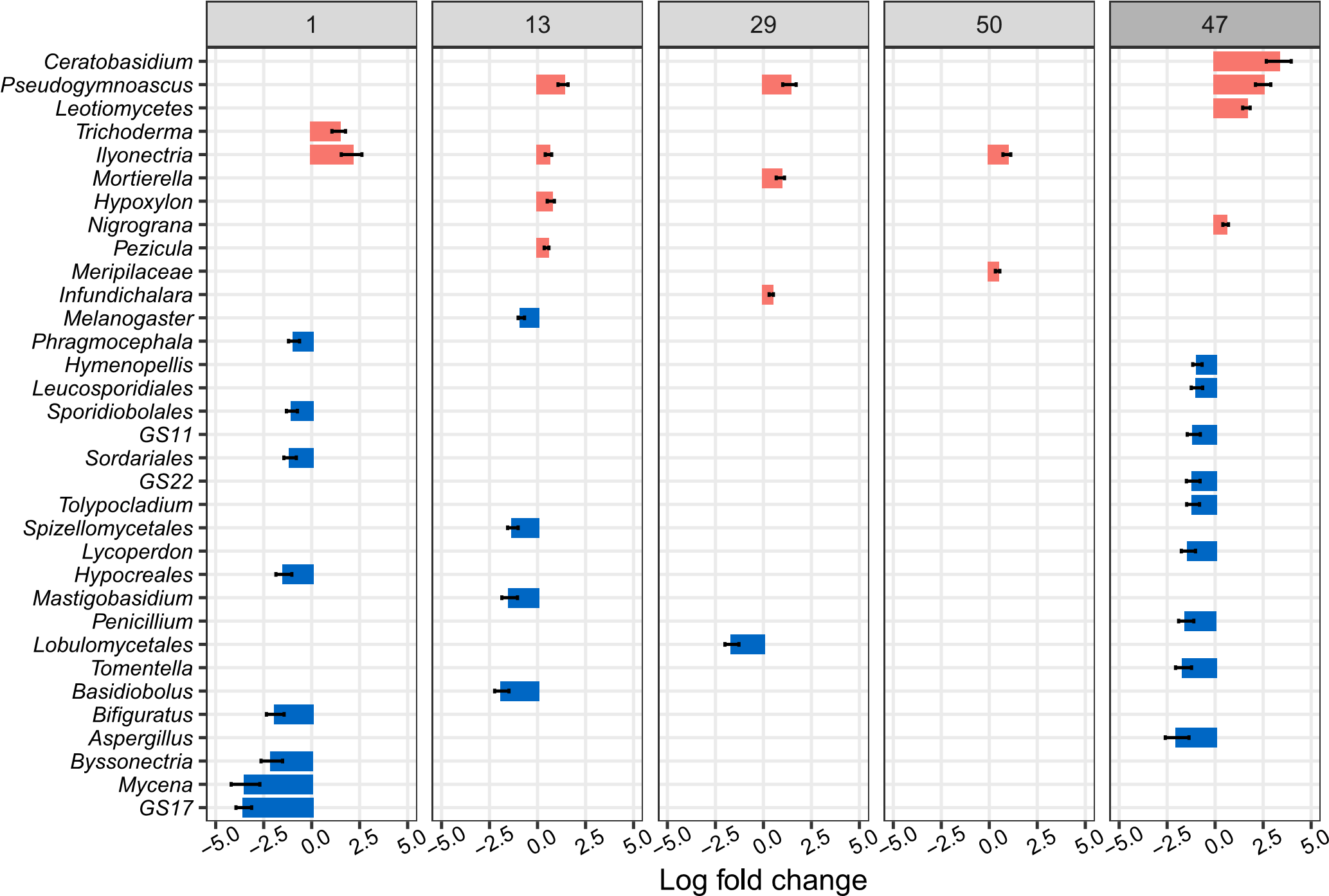

Fig. S7 Plots show fungal genera which were significantly enriched (positive log fold changes) and significantly depleted (negative log fold changes) on model cellulose particles compared to the soil, in which particles had been incubated, one, 13, 29, and 50 days after the start of the lab incubation and 47 days after the start of the field incubation. Log fold changes are calculated from relative abundance data. Bars represent the positive or negative log fold change, a measure of effect size, in the relative abundance of significantly enriched genera (FDR-adjusted P-value < 0.05). Differentially abundant genera were identified using the ANCOM-BC package (Lin and Peddada, 2020). Error bars represent the standard error of the log fold change. We controlled for false discovery rates with the Benjamini-Hochberg multiple test correction.

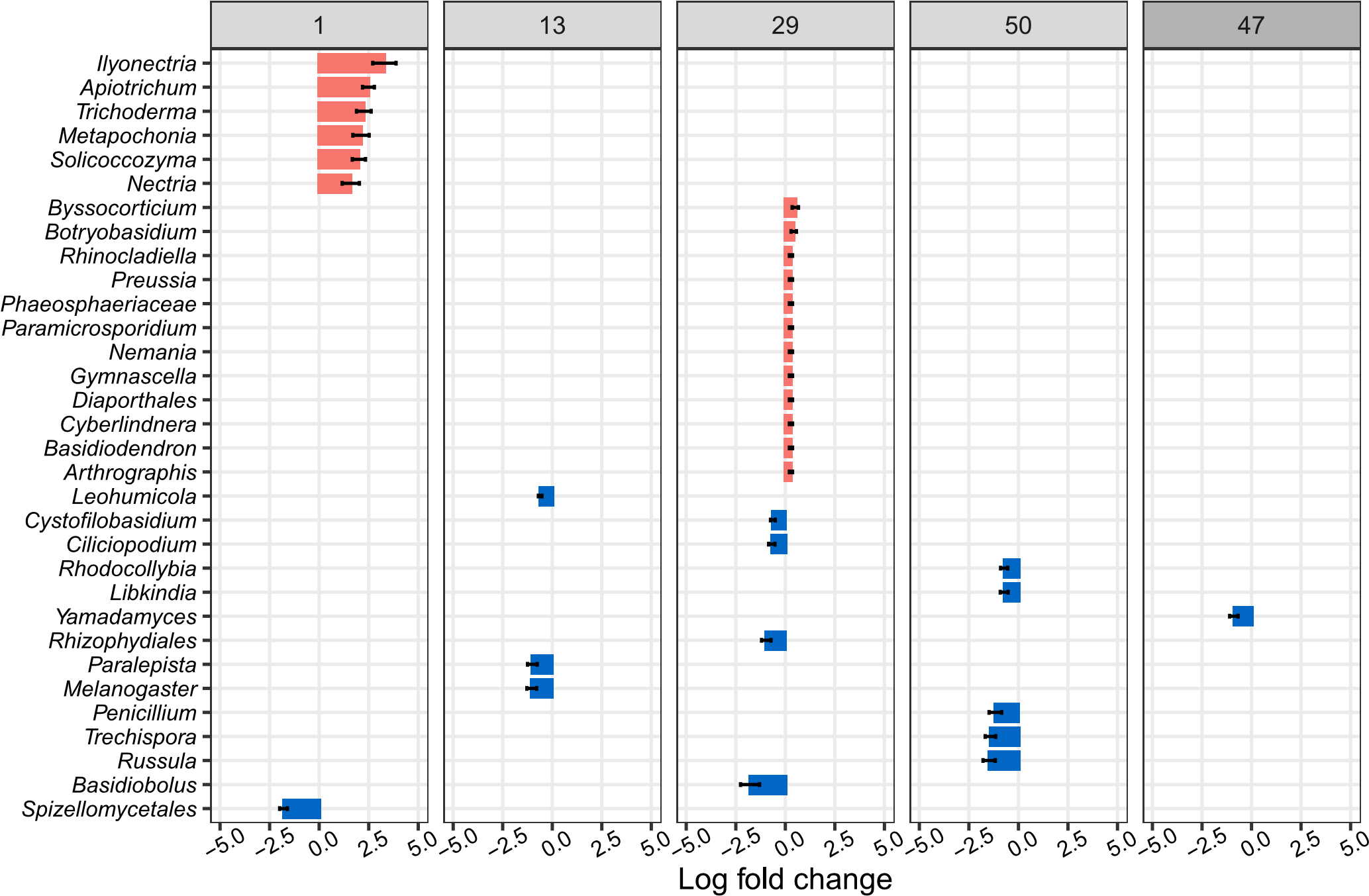
Fig. S8 Plots show fungal genera which were significantly enriched (positive log fold changes) and significantly depleted (negative log fold changes) on “no substrate” model particles compared to the soil, in which particles had been incubated, one, 13, 29, and 50 days after the start of the lab incubation and 47 days after the start of the field incubation. Log fold changes are calculated from relative abundance data. Bars represent the positive or negative log fold change, a measure of effect size, in the relative abundance of significantly enriched genera (FDR-adjusted P-value < 0.05). Differentially abundant genera were identified using the ANCOM-BC package (Lin and Peddada, 2020). Error bars represent the standard error of the log fold change. We controlled for false discovery rates with the Benjamini-Hochberg multiple test correction.

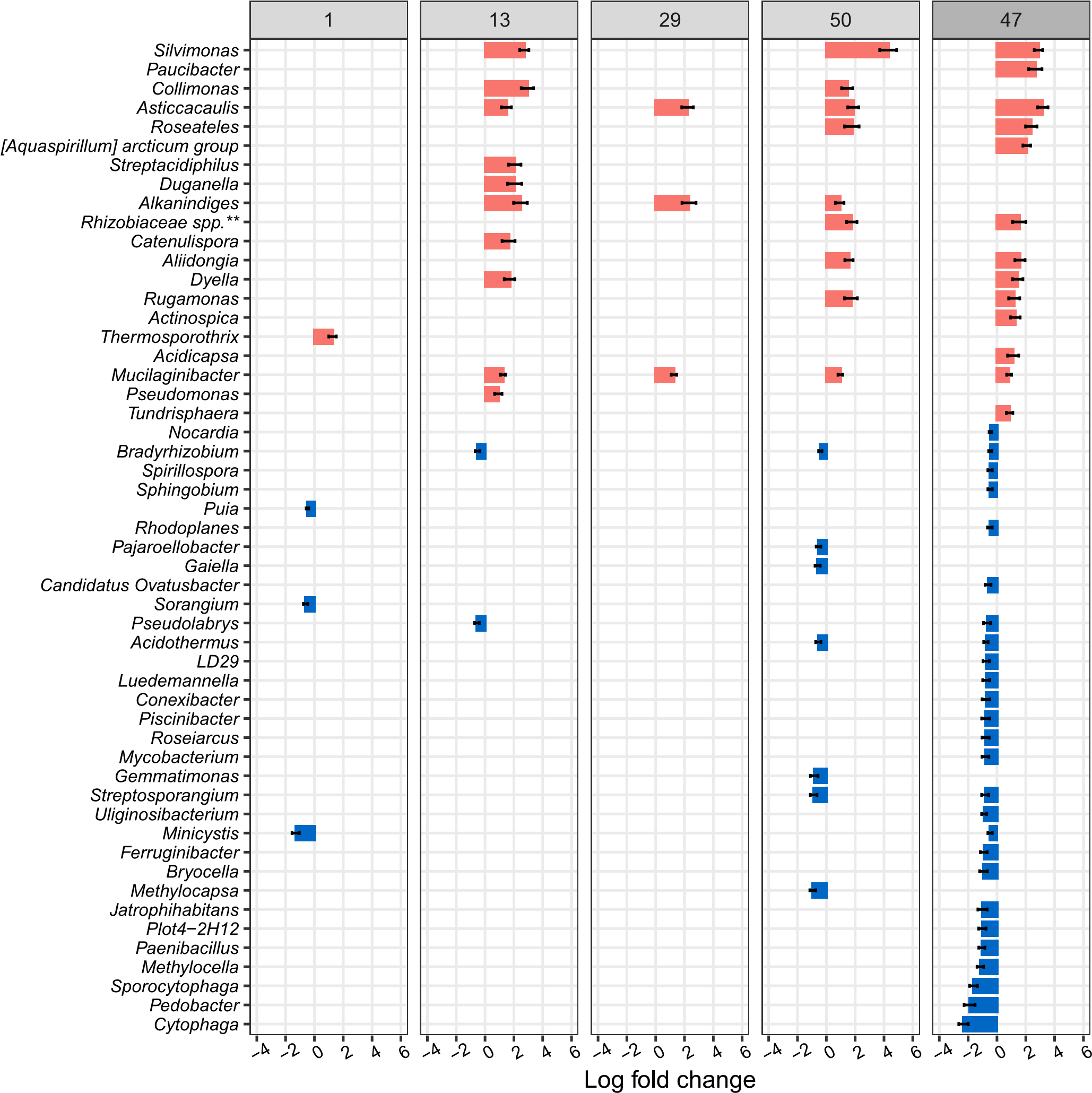

Fig. S9 Plots show bacterial genera which were significantly enriched (positive log fold changes) and significantly depleted (negative log fold changes) on chitin model particles compared to “no substrate” model particles one, 13, 29, and 50 days after the start of the lab incubation and 47 days after the start of the field incubation. Log fold changes are calculated from relative abundance data. Bars represent the positive or negative log fold change, a measure of effect size, in the relative abundance of significantly enriched genera (FDR-adjusted P-value < 0.05). Differentially abundant genera were identified using the ANCOM-BC package (Lin and Peddada, 2020). Error bars represent the standard error of the log fold change. We controlled for false discovery rates with the Benjamini-Hochberg multiple test correction. Rhizobiaceae spp.** refers to the Allorhizobium-Neorhizobium-Pararhizobium-Rhizobium clade.

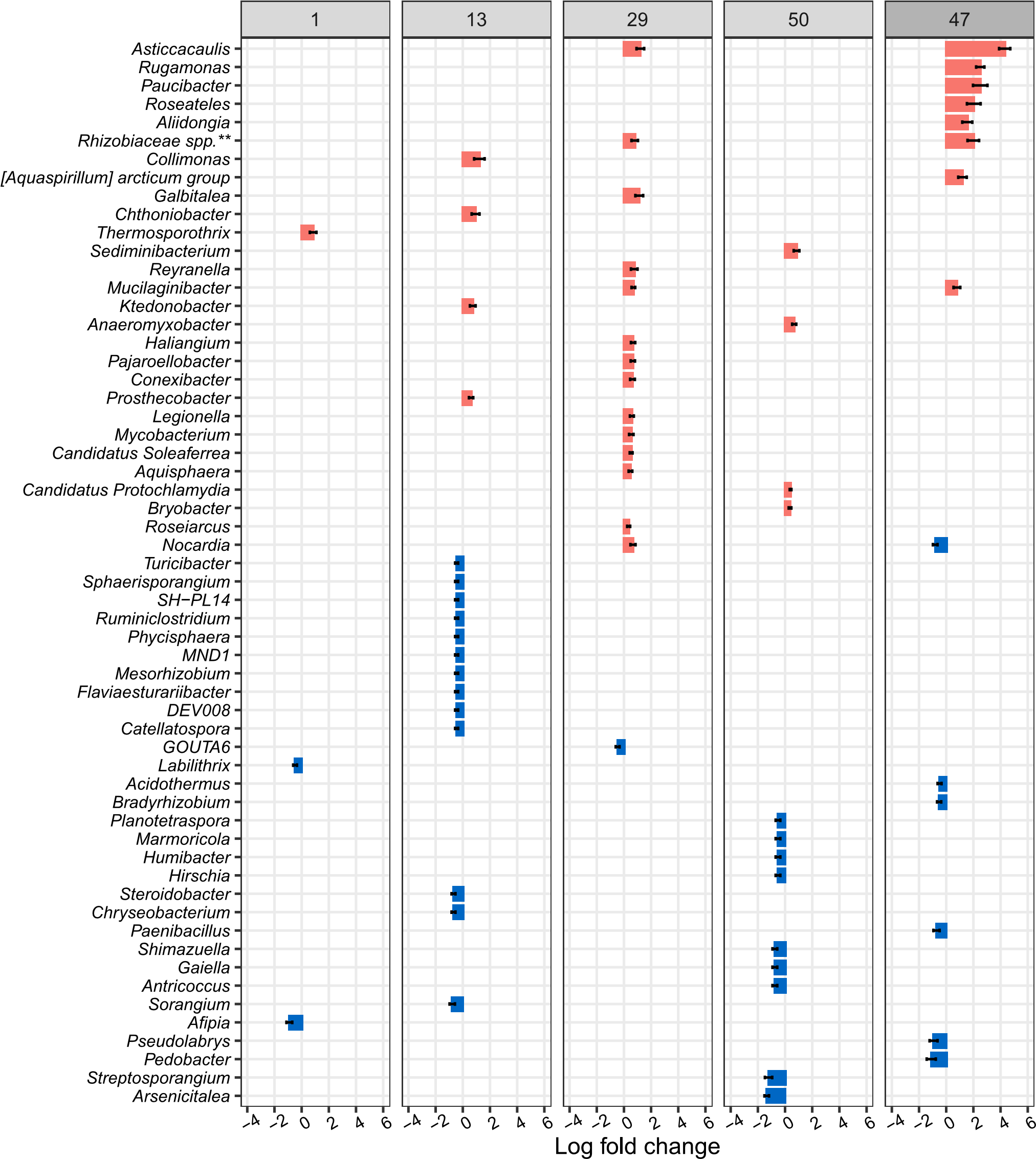

Fig. S10 Plots show bacterial genera which were significantly enriched (positive log fold changes) and significantly depleted (negative log fold changes) on cellulose model particles compared to “no substrate” model particles one, 13, 29, and 50 days after the start of the lab incubation and 47 days after the start of the field incubation. Log fold changes are calculated from relative abundance data. Bars represent the positive or negative log fold change, a measure of effect size, in the relative abundance of significantly enriched genera (FDR-adjusted P-value < 0.05). Differentially abundant genera were identified using the ANCOM-BC package (Lin and Peddada, 2020). Error bars represent the standard error of the log fold change. We controlled for false discovery rates with the Benjamini-Hochberg multiple test correction. *Rhizobiaceae spp.*** refers to the *Allorhizobium-Neorhizobium-Pararhizobium-Rhizobium* clade.

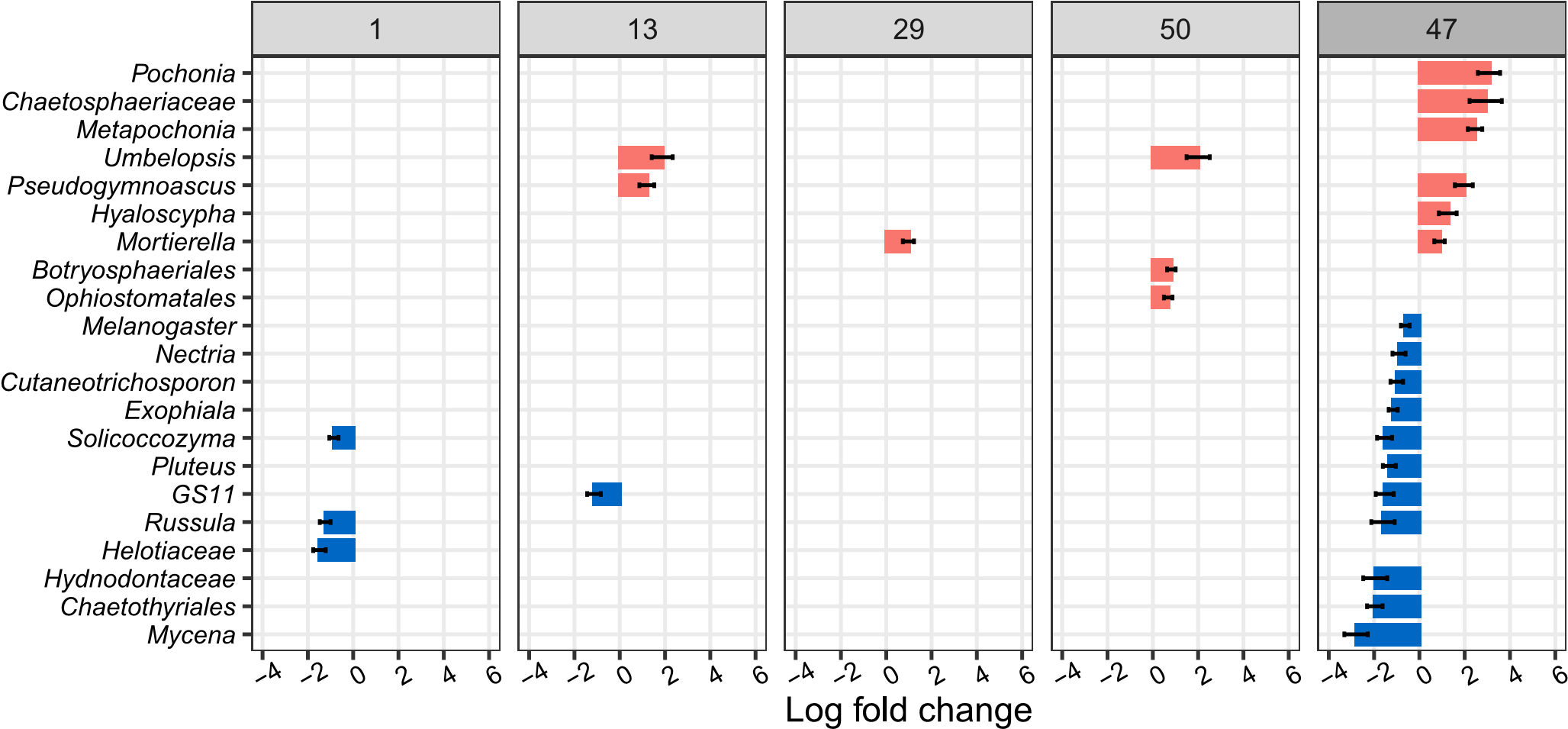

Fig. S11 Plots show fungal genera which were significantly enriched (positive log fold changes) and significantly depleted (negative log fold changes) on chitin model particles compared to “no substrate” model particles one, 13, 29, and 50 days after the start of the lab incubation and 47 days after the start of the field incubation. Log fold changes are calculated from relative abundance data. Bars represent the positive or negative log fold change, a measure of effect size, in the relative abundance of significantly enriched genera (FDR-adjusted P-value < 0.05). Differentially abundant genera were identified using the ANCOM-BC package (Lin and Peddada, 2020). Error bars represent the standard error of the log fold change. We controlled for false discovery rates with the Benjamini-Hochberg multiple test correction.

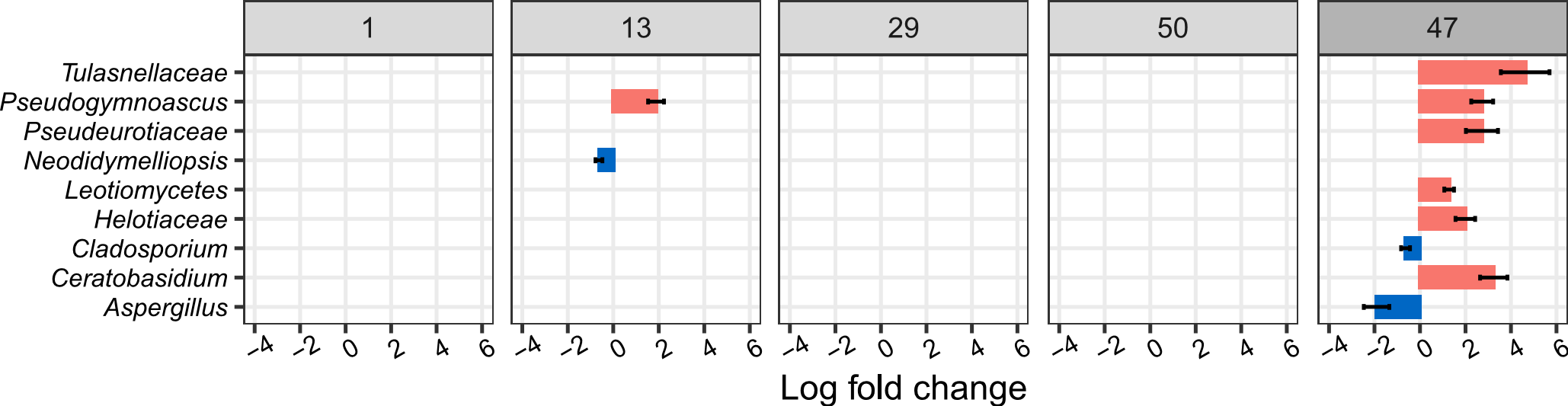

Fig. S12 Plots show fungal genera which were significantly enriched (positive log fold changes) and significantly depleted (negative log fold changes) on cellulose model particles compared to “no substrate” model particles one, 13, 29, and 50 days after the start of the lab incubation and 47 days after the start of the field incubation. Log fold changes are calculated from relative abundance data. Bars represent the positive or negative log fold change, a measure of effect size, in the relative abundance of significantly enriched genera (FDR-adjusted P-value < 0.05). Differentially abundant genera were identified using the ANCOM-BC package (Lin and Peddada, 2020). Error bars represent the standard error of the log fold change. We controlled for false discovery rates with the Benjamini-Hochberg multiple test correction.
